# Relict duck-billed dinosaurs survived into the last age of the dinosaurs in subantarctic Chile

**DOI:** 10.1101/2023.03.04.531097

**Authors:** Jhonatan Alarcón-Muñoz, Alexander O. Vargas, Hans Püschel, Sergio Soto-Acuña, Leslie Manríquez, Marcelo Leppe, Jonatan Kaluza, Verónica Milla, Carolina Simon-Gutstein, José Palma-Liberona, Wolfgang Stinnesbeck, Eberhard Frey, Juan Pablo Pino, Dániel Bajor, Elaine Núñez, Héctor Ortiz, Héctor Mansilla, David Rubilar-Rogers, Penélope Cruzado-Caballero

## Abstract

In the dusk of the dinosaur era, the advanced duck-billed dinosaurs (Family Hadrosauridae) are thought to have outcompeted other herbivores, making ecosystems less diverse and more vulnerable to the Cretaceous-Paleogene asteroid impact. They were also among the first terrestrial organisms to disperse from North America into South America. Here, we present the first new species of subantarctic duck-billed dinosaur, CPAP 3054, of early Maastrichtian age in Magallanes, Chile. Surprisingly, unlike duckbills further north in Patagonia, CPAP 3054 is not an advanced duckbill, but descends from North American forms that were transitional to Hadrosauridae, diverging shortly before the origin of this family. In North America, these forms were replaced by hadrosaurids in the late Campanian. The survival into the Maastrichtian of a pre-hadrosaurid lineage suggests the ancestors of CPAP 3054 arrived earlier in South America than the hadrosaurids, reaching further south before the Cretaceous-Paleogene mass extinction, where they avoided competition from hadrosaurids.

**Additional note:** This work contains a new biological name. New names in preprints are not considered available by the ICZN. To avoid ambiguity, the new biological name is not included in this preprint, and the holotype specimen number CPAP 3054 is used as a placeholder. Paratypes described in this preprint are also used in the diagnosis.

## Introduction

Duck-billed dinosaurs (Hadrosauroidea) are among the most successful herbivores in the history of life on earth. Their tooth batteries with hundreds of teeth are arguably the most complex in vertebrate evolution (1), and were capable of crushing, grinding, and shearing, allowing them to exploit a broad range of plant resources that they could further reach at both short and high distances above ground (2). The duck-billed dinosaurs attained their maximum historical diversity upon the radiation of advanced forms of the family Hadrosauridae (the “true” duckbills), during the Campanian-Maastrichtian. In this time period, they are thought to have outcompeted other herbivores, contributing to overall declines of dinosaur biodiversity in North America and Central China previous to the asteroid impact of the Cretaceous-Paleogene (K-Pg) mass extinction (2). Also in this time period, Hadrosaurids were among the very few dinosaurs from Laurasia that are known to have colonized Gondwanan continents (3, 4). One species has been described from northern Africa (5) and 5 from South America, all within central and northern Patagonia (Chubut and Neuquén) (4, 6, 7,8, 9). There is also evidence that duck-billed dinosaurs reached further into southern Patagonia, and even the Antarctic peninsula (10,11,12). These regions at the time retained close geographic proximity with intermittent formation of land bridges and intercontinental biotic exchanges (13); further, they may have been at least partially isolated from the rest of South America due to marine transgressions in the latest Cretaceous such as the Kawas Sea (14,15,16,17). Dinosaur faunas in these far southern regions are poorly known, and duck-billed dinosaurs are only documented by few and partial unnamed remains (11,12,13), which are currently assumed to belong to hadrosaurids like those that inhabited central and north Patagonia. Although Antarctica remained close to Australia and New Zealand, and often exchanged biotic components (10), the absence of any confirmed remains of duck-billed dinosaurs in the latter two suggests duckbills did not have enough time to arrive there before the K-Pg mass extinction.

In 2013, we initiated excavation of a monodominant bonebed with abundant disarticulated remains from several individuals of a new species of subantarctic duck-billed dinosaur (18, 19). The site is of early Maastrichtian age (see Methods) and is located in the Río de Las Chinas valley of the Magallanes Region of subantarctic Chile, at the southernmost tip of southern Patagonia (51°S, Fig. 1). Here, we will use the term “subantarctic” in the physiographic sense, to refer to the territories that are immediately north of the Antarctic convergence, being roughly south of 46°S (the southern limit of Chubut, central Patagonia). So far, excavation of about 28 m2 of a layer approximately 80 cm thick has yielded over 42 skeletal elements, but other unexcavated outcrops of the bonebed are found in an extension of about 5 km (see Methods). The materials recovered so far are the most informative for any duck-billed dinosaur this far south, yielding unexpected conclusions. Instead of being specifically related to other South American duckbills, the new dinosaur is not a hadrosaurid: instead, it is similar to North American forms that diverged shortly before the origin of that family. In technical terms, it is a non-hadrosaurid hadrosauroid that is closely related to Hadrosauridae. For the sake of simplicity, we will refer to it as a “transitional” duckbill, to emphasize that its morphology is transitional to that of the “true” duckbills of the family Hadrosauridae.

**Fig. 1.**
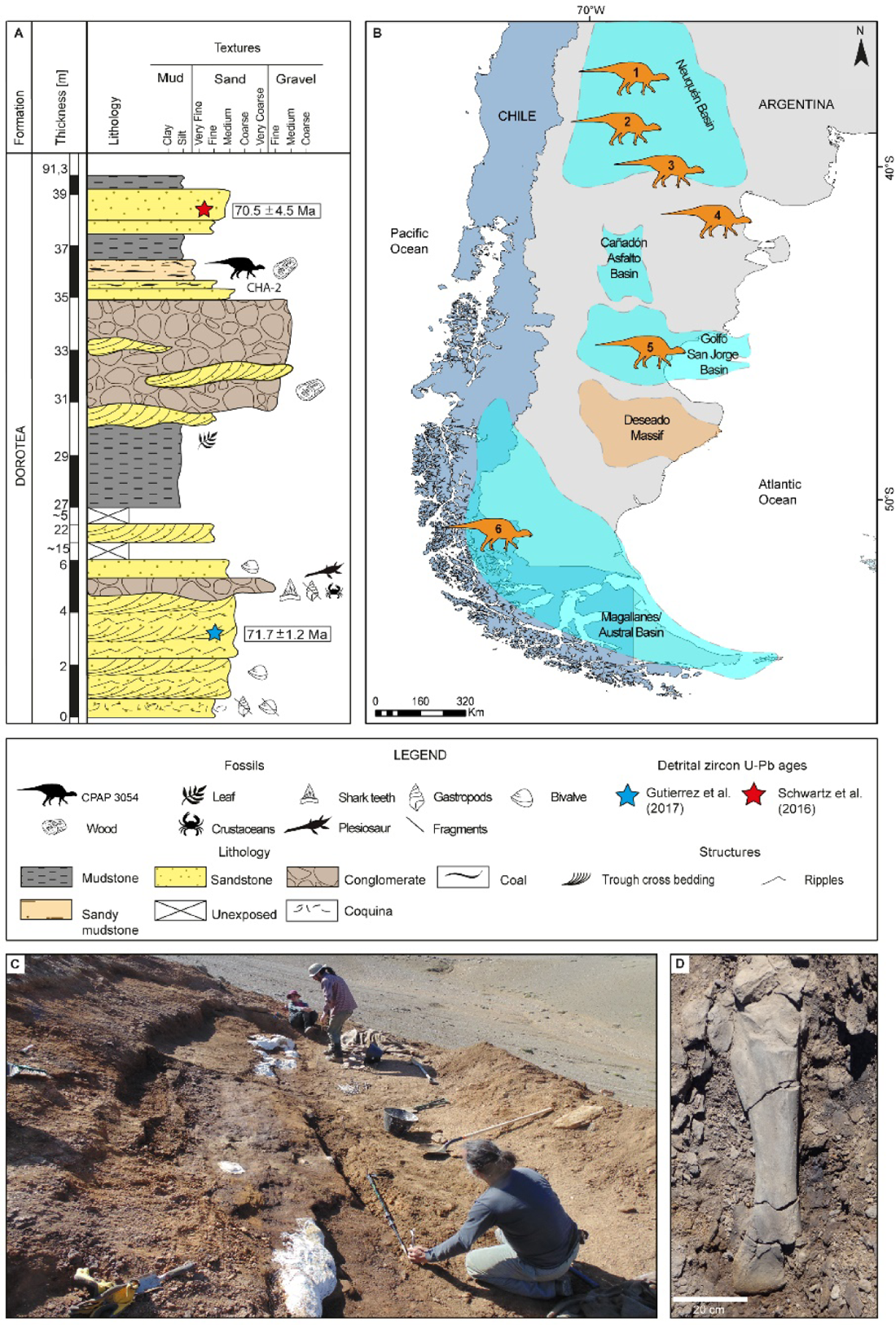
Stratigraphic location of CPAP 3054 and geographic distribution of South American duck-billed dinosaurs. **A**, Stratigraphic section of the Dorotea Formation, in which the location of the bones of CPAP 3054 is indicated. **B**, Location of the duck-billed dinosaurs from South America: **1**, *Lapampasaurus cholinoi* (Islas Malvinas locality, La Pampa Province, late Campanian-early Maastrichtian, Allen Formation); **2**, *Kelumapusaura machi* (Matadero Hill, General Roca city, Río Negro Province, middle Campanian-early Maastrichtian, Allen Formation); **3**, *Bonapartesaurus rionegrensis* (Salitral Moreno, General Roca Department, Río Negro Province, middleCampanian-early Maastrichtian, Allen Formation); **4**, *Huallasaurus australis* (Arroyo Verde Puelén Departament, Río Negro Province, late Campanian-early Maastrichtian, Los Alamitos Formation); **5**, *Secernosaurus koerneri* (Río Chico, east of Lake Colhué Huapi, Chubut Province, Maastrichtian, Lago Colhué Huapi Formation); **6**, CPAP 3054 (Río de las Chinas Valley, Magallanes Region, late Campanian-early Maastrichtian, Dorotea Formation). **C**, The quarry from which CPAP 3054 bones were excavated (bonebed level); **D**, detail of a nearly complete tibia.

## Results

### Systematic Paleontology

Dinosauria Owen, 1842

Ornithischia Seeley, 1887

Ornithopoda Marsh, 1881

Hadrosauroidea Cope, 1870 CPAP 3054 (gen. et. sp. nov.)

### Holotype

CPAP 3054, right ilium.

CPAP 3054 was discovered among several other bones; all skeletal elements were disarticulated, precluding a confident assignment of multiple elements to specific individuals. We therefore designated CPAP 3054 as the holotype because of evident diagnostic features. The preacetabular process of CPAP 3054 is incomplete but shows a “T” shaped cross-section due to the well-developed lateral and medial crests that characterize adults (20, 21). These features allowed us to discard a juvenile status (that is, an animal with no signs of impending maturity, 22).

### Paratypes

Several disarticulated bones were selected as paratypes of the same species (Fig. 2 and Figs. S1-S3) given the lack of evidence for more than one species at the site, and the fact that monodominant associations are common in the fossil record of duck-billed dinosaurs, reflecting social interaction or herding behavior (23,24,25,26,27).

**Fig. 2.**
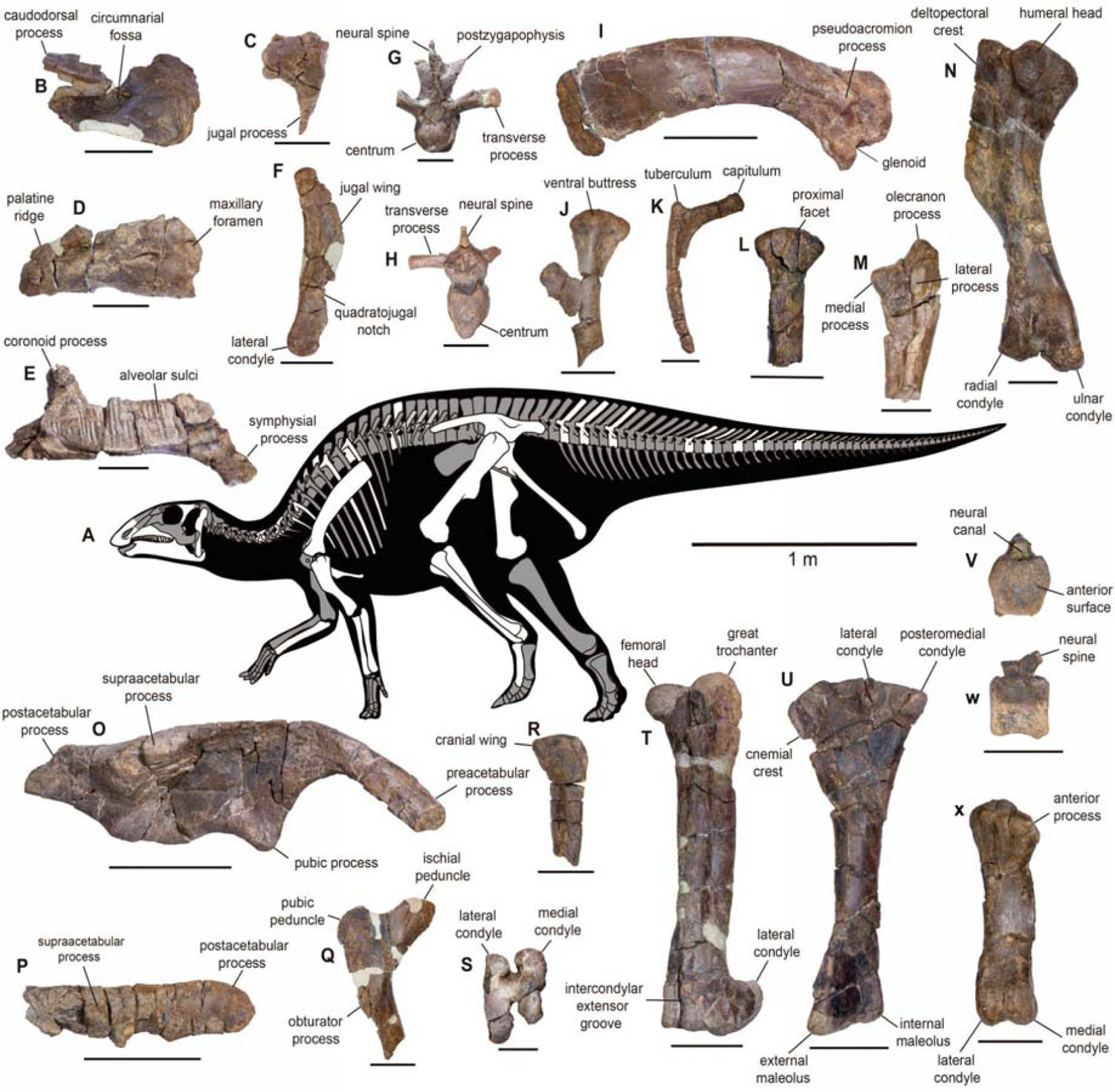
CPAP 3054, skeletal anatomy. **A**, Bones described in this work (white). Some elements are indicated specularly to facilitate their representation. **B**, **CPAP 5337**, right premaxilla in lateral view. **C**, **CPAP 5341**, incomplete left postorbital in lateral view. **D**, **CPAP 5340**, incomplete right maxilla in lateral view. **E**, **CPAP 5370**, left dentary in medial view. **F**, **CPAP 5343**, left quadrate in lateral view. **G**, **CPAP 5344**, cervical vertebra in anterior view. **H**, **CPAP 5345**, dorsal vertebra in anterior view. **I**, **CPAP 5371**, right scapula in lateral view. **J**, **CPAP 5352**, left sternum in ventral view. **K**, **CPAP 5378**, incomplete right rib in anterior view. **L**, **CPAP 5379**, proximal portion of right radius in posterior view. **M**, **CPAP 5355**, incomplete left ulna in anterolateral view. **N**, **CPAP 5353**, left humerus in lateral view. **O**, **CPAP 3054** (holotype), right ilium in lateral view. **P**, **CPAP 5356**, left postacetabular process in lateral view. **Q**, **CPAP 5357**, proximal portion of left isquion in lateral view. **R**, **CPAP 5363**, proximal portion of right fibula in lateral view. **S**, **CPAP 5360**, incomplete right femur in distal view. **T**, **CPAP 5358**, left femur in anterior view. **U**, **CPAP 5362**, left tibia in lateral view. **V**, **W**, **CPAP 5349**, caudal vertebra in anterior and lateral views. **X**, **CPAP 5364**, right metatarsal III in anterior view. Scale bars, B-G, J, K-N, Q, S, V-X= 50 mm; H-I, O-P, R, T-U=100 mm.

Taphonomic aspects such as Hydraulic equivalence and lack of abrasion also suggest that bones at the site underwent minimal transport, discarding water currents as the reason for accumulated remains of numerous individuals (see Taphonomy section in Supplementary Materials and Tables S1 and S2). The bones assigned as paratypes are the following: a right premaxilla (CPAP 5337), the incomplete left maxilla of a juvenile individual (CPAP 5338), an incomplete left maxilla (CPAP 5339), an incomplete right maxilla (CPAP 5340), an incomplete left postorbital (CPAP 5341), an incomplete right dentary (CPAP 5342), a left dentary (CPAP 5370), a fragmentary right dentary (CPAP 5368), a right quadrate (CPAP 5343), a cervical vertebra (CPAP 5344), a cervical centrum (CPAP 5380), two incomplete dorsal vertebrae (CPAP 5345 and CPAP 5346), a dorsal neural arch (CPAP 5377), eight proximal and mid caudal vertebrae (CPAP 5347, CPAP 5348, CPAP 5349, CPAP 5350, CPAP 5351, CPAP 5384, CPAP 5385 and CPAP 5386), four incomplete ribs (CPAP 5378, CPAP 5381, CPAP 5382 and CPAP 5383), a right scapula (CPAP 5371), an incomplete left sternum (CPAP 5352), a complete left humerus (CPAP 5353), an incomplete left humerus (CPAP 5354), two fragments of a left humerus (CPAP 5369), an incomplete left ulna (CPAP 5355), an incomplete right ulna (CPAP 5373), an incomplete right radius (CPAP 5379), a postacetabular process with part of the iliac blade (CPAP 5356), an incomplete right ischium (CPAP 5357), a complete left femur (CPAP 5358), an incomplete right femur (CPAP 5359), the distal end of a right femur (CPAP 5360), the distal end of the right femur of a juvenile individual (CPAP 5361), a complete left tibia (CPAP 5362), an incomplete left tibia (CPAP 5372), an incomplete right fibula (CPAP 5363), a right metatarsal III (CPAP 5364).

### Etymology

The words used for the new species are in the language of the Aónik’enk, the indigenous people that inhabited the region where the new species was discovered (28).

### Type locality and horizon

El Puesto Area (50°42’42“S y 72°32’29” W), Río de las Chinas Valley, Estancia Cerro Guido, Magallanes Region, Chilean Patagonia (51° S). Upper section of the Dorotea Formation (lower Maastrichtian), between 71.7 ± 1.2 and 70.5 ± 4.5 million years (29, 30) (See Methods).

### Diagnosis

Small-sized hadrosauroid dinosaur (total body length ∼ 4 m) that differs from all other members of this clade in possessing a scapula with an antero-ventrally curved pseudoacromion process, and an ilium with a medioventrally and anteroposteriorly well-developed sacral crest extending posterior to the base of the postacetabular process. CPAP 3054 also possesses a unique combination of characters, differing from Hadrosauridae in a maxilla with a jugal articular surface showing a prominent and caudally projecting dorsal tubercle; a dentary with a short diastema, mandibular symphysis oblique relative to its long axis, and tooth rows with less than 30 tooth positions, that do not extend beyond the coronoid process, and converge anteriorly with the lateral surface of the dentary; quadrate with a medial condyle that is not markedly elevated dorsally compared with the lateral condyle; a deltopectoral crest with a rounded laterodistal end extending less than 48% of the total length of the humerus; ilium with nearly straight dorsal border, and a supraacetabular crest longer than 70% of the length of the iliac blade; and tibia with a cnemial crest extending less than 50% of the total length of the bone. In addition, CPAP 3054 differs from South American Hadrosauridae by a scapula with a mediolaterally narrow coracoidal facet, with a more ventrally curved pseudoacromion process (Fig. S4), and an ilium with a more laterally curved preacetabular process, a less laterally developed supraacetabular process, and a shorter and ventrally oriented pubic peduncle (Fig. S5). CPAP 3054 differs from early-diverging Hadrosauroidea in that it presents derived characters also found in Hadrosauridae: a premaxilla with a “double-layer” morphology; a supraacetabular process anteriorly located with respect to the dorsal tuberculum of the ischial peduncle; the ratio between the base of the preacetabular process and the distance between the dorsal border of the ilium and the pubic peduncle is greater than 0.5; and a laterocaudal condyle of the proximal end of the tibia is more robust than the lateral one. Regarding similar non-hadrosaurids closely related to Hadrosauridae, CPAP 3054 differs from *Eotrachodon* in that the precircumnarial region of the premaxilla is more anteroposteriorly extended, lacking a caudally everted oral margin; and the maxilla has an articular surface for the jugal located anterior to the dorsal process rather than posterior to it. CPAP 3054 differs from *Lophorhothon* in that the jugal process of the postorbital is pointing ventrally and the posterior edge of this process is slightly convex, rather than concave; and the ilium is higher, with a laterally less developed supraacetabular crest, located anterior to the dorsal tubercle of the ischial peduncle. CPAP 3054 differs from *Huehuecanauhtlus* in that the preacetabular process of the ilium in dorsal view is markedly curved laterally, less curved ventrally, and its base is dorsoventrally narrower; the dorsal tubercle of the ischial peduncle of the ilium is less dorsally elevated; and the postacetabular process is approximately rectangular in lateral view, rather than triangular.

### Diagnosis of the genus

As for the type species.

### Description

The premaxilla has an oval, very deep circumnarial fossa which lacks accessory fossae, foramina, and ridges, unlike Saurolophinae (31,32,33,34,35,36,37, Fig.3). The oral margin is convex and anteroventrally deflected, unlike most Saurolophinae (38, 39, Fig. 3). It shows a denticulate margin, separated by a shallow groove from a denticulate border with the palatine (“double layer”), as in Saurolophidae and some non-hadrosaurid hadrosauroids (31,37,40,41,42; Fig. 4). The maxilla is subtriangular in lateral view. The articular surface for the jugal is robust, subquadrangular in shape and has a prominent dorsal tubercle, unlike hadrosaurids (43, 44). A large rostral maxillary foramen is present at the lateral side, as in Saurolophinae (31,40, Fig. 4). The medial surface bears anteroposteriorly aligned neurovascular foramina (31, 40). The teeth are poorly preserved, making it difficult to determine their morphology and number in the maxilla. The postorbital is T-shaped in lateral view. Unlike most hadrosaurids, the dentary has a short diastema (less than 20% of the length of the alveolar row), and the mandibular symphysis is oblique relative to the long axis of the dentary, as in several non-hadrosaurid duckbills (35,45,46, Fig. 3). The alveolar furrows are parallel to each other and dorsoventrally extended. The most complete dentary (CPAP 5370) shows about 25 tooth positions, rather than over 30 as in Hadrosauridae (39,40,47,48, Fig. 3). The dentary tooth row extends further posteriorly than in basal iguanodontians, reaching the coronoid process, but not extending beyond it, unlike hadrosaurids (40,44,48, Fig. 3). The tooth row converges anteriorly with the lateral surface of the dentary, unlike *Plesiohadros* and Hadrosauridae (37,40,49,50, Fig. 3). The coronoid process is robust, almost straight, and separated from the alveolar row by a groove.

**Fig. 3.**
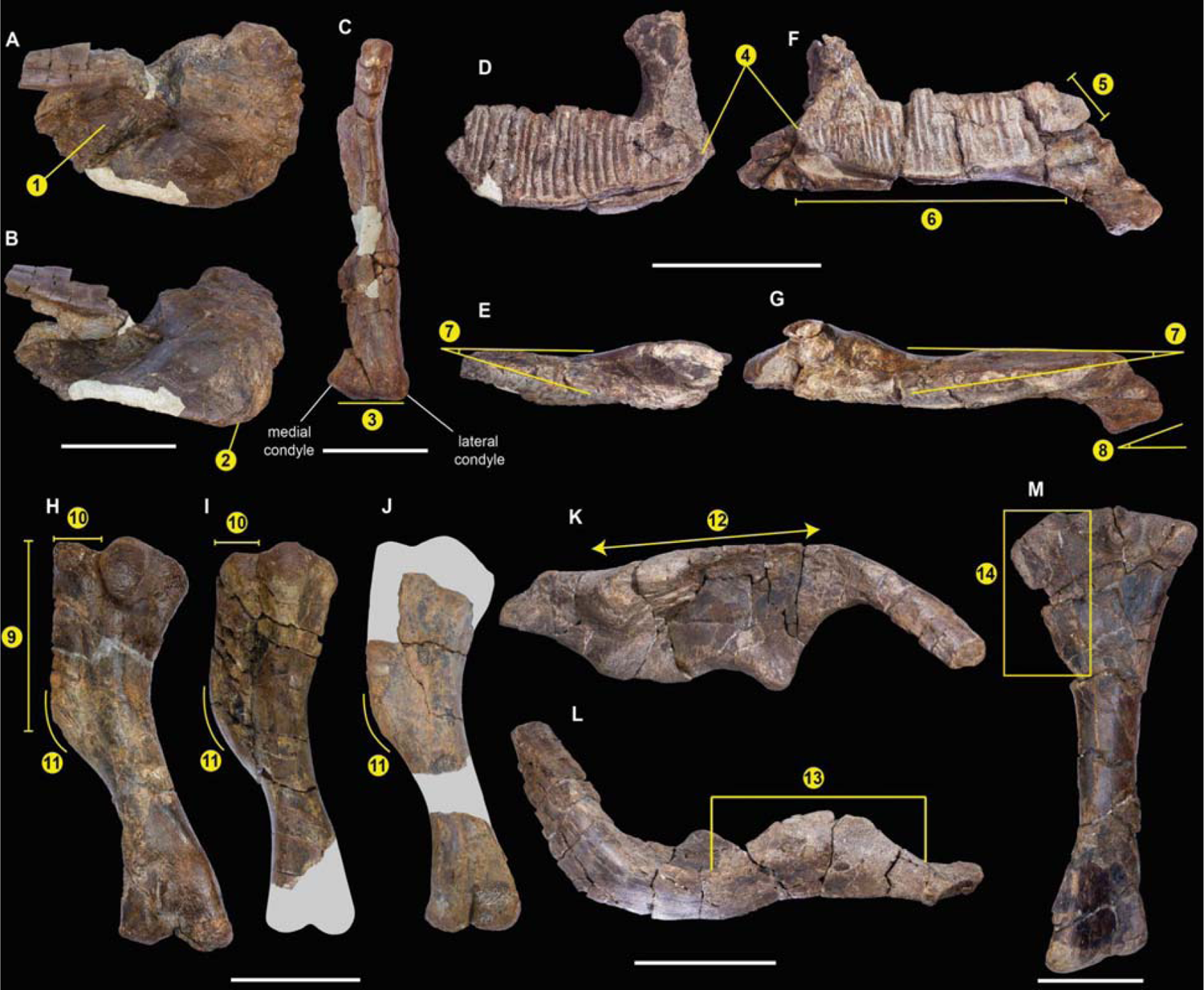
CPAP 3054 lacks several derived characters found in Hadrosauridae. A selection of plesiomorphic characters is shown. **A**-**B**, **CPAP 5337**, right premaxilla in dorsal (**A**) and lateral (**B**) views showing a circumnarial fossa without accessory fossa or foramina (1), and an anteroventrally deflected oral margin (2). **C**, **CPAP 5343**, right quadrate in posterior view. The lateral condyle is not markedly offset ventrally with respect to the medial condyle, unlike Hadrosauridae. **CPAP 5342,** right dentary in medial (**D**) and dorsal (**E**) views, and **CPAP 5370**, left dentary in medial (**F**) and dorsal (**G**) views. Tooth rows do not exceed the coronoid process (4), the diastema is short (5), there are less than 30 dental positions (6), tooth row converges anteriorly with the lateral surface of the dentary (7), and the symphysis is oblique (8). **CPAP 5353** (**H**), **CPAP 5354** (**I**), and **CPAP 5369** (**J**), left humeri in posterolateral view, in which the ratio between length of the deltopectoral crest and the total length of the humerus is lower than 0.48 (9), and the deltopectoral crest is mediolaterally short (10), with a widely arcuate laterodistal corner (11). **K**-**L**, **CPAP 3054**, right ilium in lateral (**K**) and dorsal (**L**) views. The dorsal border of the ilium is almost straight (12), and the length of the supraacetabular crest is higher than 70% of the length of the iliac blade (13). **M**, **CPAP 5362**, left tibia in lateral view. In CPAP 3054, the length of the cnemial crest is lower than the 50% of the total length of the bone (14). Scale bars=50 mm (**A**-**C**) and 100 mm (**D**-**M**).

**Fig. 4.**
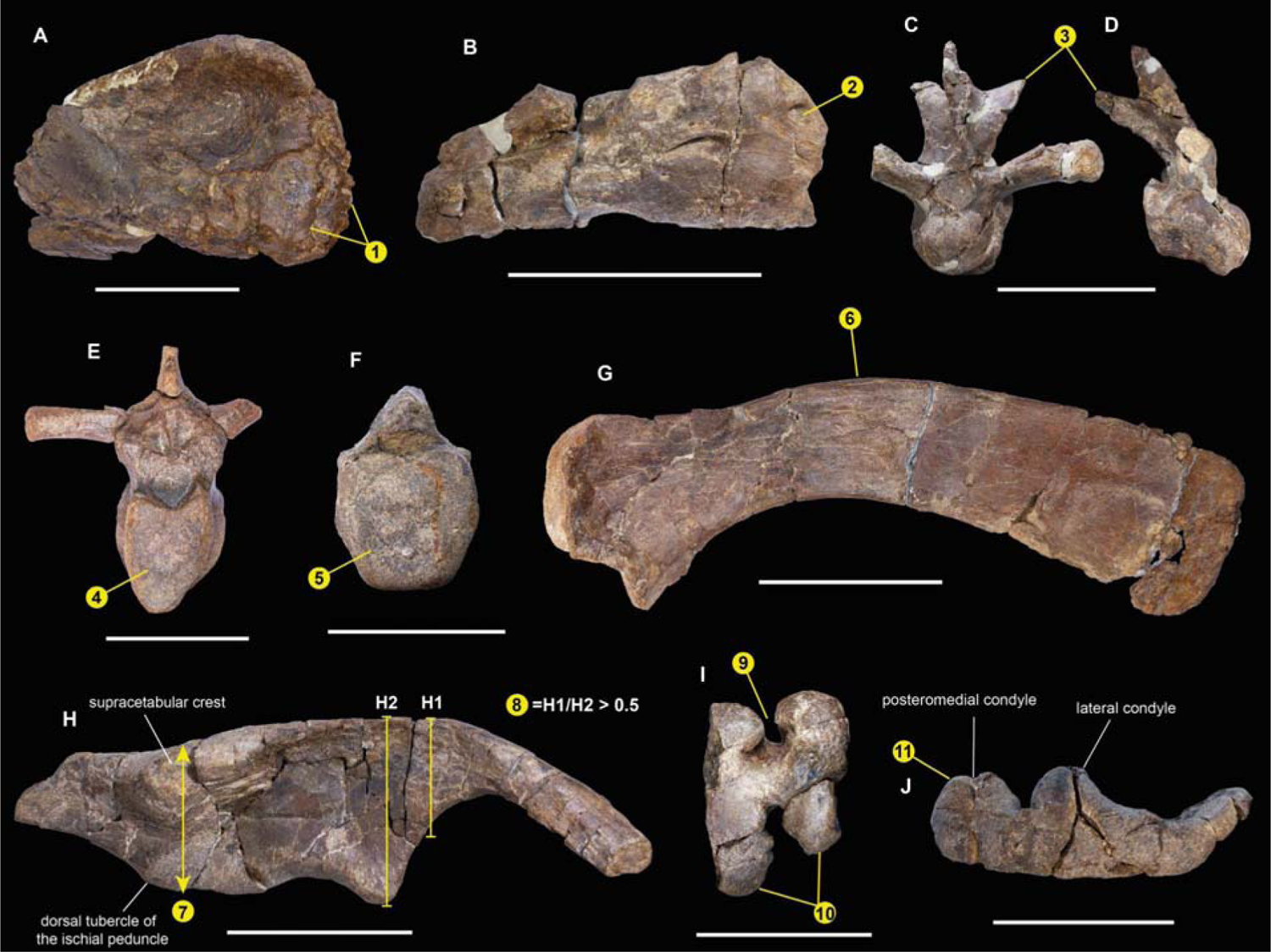
Derived characters of CPAP 3054 shared with Hadrosauridae. *G.nanoi* supports these characters evolved previous to the origin of this family. **A**, **CPAP 5337**, right premaxilla in ventral view, which shows a “double-layer” morphology (1). **B**, **CPAP 5340**, right maxilla in lateral view, with a rostral maxillary foramen on the lateral surface of the bone (2). **C**-**D**, **CPAP 5344**, mid-cervical vertebra in anterior (**C**), and lateral view (**D**). This cervical vertebra has elevated and well developed postzygapophyseal processes, which are dorsally arched (3). **E**, **CPAP 5346**, dorsal vertebra whose centrum has a “heart-shaped”articular surface (4). **F**, **CPAP 5347**, mid-caudal vertebra with a hexagonal articular surface (5). **G**, **CPAP 5371**, right scapula in medial view, which is dorsally arched (6). **H**, **CPAP 3054**, right ilium in lateral view. The ratio between the base of the preacetabular process (H1) and the distance between the dorsal border of the ilium and the pubic peduncle (H2) is greater than 0.5 (8). Furthermore, the supraacetabular process is anteriorly located with respect to the dorsal tuberculum of the ischial peduncle. **I**, **CPAP 5360**, distal end of the right femur in distal view, showing a deep intercondylar extensor groove, in which the condyles nearly meet anteriorly (9). In addition, the condyles are posteriorly well-developed (10). **J**, **CPAP 5362**, left tibia in proximal view, in which the posteromedial condyle is wider than the lateral condyle. Scale bars=50 mm (**A**, **F**), and 100 mm (**B**-**E**, **G**-**J**).

The quadrate is dorsoventrally elongated. Its dorsal half curves slightly posteriorly and lacks a squamosal buttress. The quadratojugal notch is ventrally positioned. Unlike most hadrosaurids, the lateral condyle is not markedly offset ventrally with respect to the medial condyle (40, 45, Fig. 3). The cervical centrum is longer than dorsoventrally high, and strongly opisthocoelous. The anterior and posterior articular surfaces are oval. The postzygapophyses are well elevated dorsally to the roof of the neural canal and curved posteriorly and laterally, which is common in hadrosaurids (31; Fig. 4). The dorsal centra are elongate, higher than wide, opisthocoelous and have heart-shaped articular faces like hadrosaurids and at least some non-hadrosaurid hadrosauroids (31,33,42, Fig. 4). The caudal centra are amphiplatyan with hexagonal articular contours, which is typical but not exclusive to hadrosaurids (31,33,39,51 Fig. 4).

The sternum is hatchet-shaped. The scapula has a dorsally curved blade as is typical of hadrosaurids (40; Fig. 4), with a rectangular posterior end. The deltoid ridge is poorly developed. The glenoid fossa narrows mediolaterally and the pseudoacromion process is prominent and anteroventrally curved, unlike in all other South American duckbills (Fig. S4). In addition, unlike South American hadrosaurids such as *Huallasaurus*, *Lapampasaurus* and *Kelumapusaura*, the coracoidal facet is mediolaterally narrower (Fig. S4).

Unlike Hadrosauridae, the deltopectoral crest of the humerus extends over 45% the total length of the bone (40, 52). The ratio between length and maximum width of the deltopectoral crest is 3.9, which is higher than in Saurolophidae (20). The laterodistal corner of the deltopectoral crest is widely arcuate, unlike most hadrosaurids (40,52,53, Fig. 3). The ulna has a prominent olecranon process. The iliac peduncle of the ischium is longer than the pubic peduncle. The dorsal border of the ilium is almost straight, unlike *Huehuecanauhtlus* and hadrosaurids, including South American duckbills (33,35,39,52,54,55,56, Fig. S5). The sacral crest is prominent and more developed than in other South American duckbills (Fig. S5). The preacetabular process is incomplete. This is robust, with a “T” shaped cross-section, and is more laterally curved than in other South American duckbills (Fig. S5). The preserved anterior end of the preacetabular process surpasses the lateral extension of the supraacetabular process, as in *Secernosaurus*, but not in other South American duckbills (Fig. S5). The ratio between the maximum height of the caudal end of the preacetabular process and the distance between the dorsal margin of the ilium and the ventral edge of the pubic peduncle is 0.58, which is higher than in non-hadrosaurid hadrosauroids (9,40, Fig. 4). The pubic peduncle is robust and triangular in lateral view, unlike the elongated, rod-like pubic peduncle of basal iguanodontoids such as *Iguanodon* (57). The pubic peduncle is ventrally oriented and shorter than in some South American duckbills (Fig. S5). The ischial peduncle consists of two low and blunt tubercles of similar size. The supraacetabular process projects slightly lateroventrally, with its ventral apex slightly anteriorly located with respect to the dorsal tubercle of the ischial peduncle, as in Hadrosauridae (20, 52, Fig. 4). The length of the supraacetabular crest reaches approximately 82.6 % of the length of the central plate of the ilium. This is within the range of non-hadrosaurids: lower than in basal iguanodontoids such as *Iguanodon* (>85%), and higher than in Hadrosauridae (<70%) (50, Fig. 3). The lateroventral protrusion of the supraacetabular process is less marked than in other South American duckbills except for *Secernosaurus* (Fig. S5). The postacetabular process is rectangular, slightly dorsally inclined, and twisted. The femur is straight, with a triangular fourth trochanter. The flexor and extensor grooves are deep, with the latter almost closed, and the distal condyles are strongly projected posteriorly, which are features typical of hadrosaurids (31, Fig. 4). The lesser trochanter is small and well-separated from the greater trochanter. The cnemial crest extends to less than half the total length of the bone, which differs from all hadrosaurids except *Bonapartesaurus* (9,45,56,58,59, Fig. 3). Posterior to the cnemial crest there are two rounded condyles separated by a sulcus. The posteromedial condyle is more robust and longer than the lateral condyle, like Hadrosauridae (60, Fig. 4). The tibia has a torsion of approximately 45° between the proximal and distal epiphysis. The lateral malleolus is more extended distally than the medial one. A fragment of a fibula reveals an enlarged proximal end. As in several non-hadrosaurids, the articular surface of the metatarsal III is triangular (33, 58); the ratio between the length and mediolateral width of this bone is 4.5.

## Discussion

CPAP 3054 is a duck-billed dinosaur that shares some derived traits with “true” duckbills of the family Hadrosauridae (Fig. 4), while several other characters retain their ancestral condition (Fig. 3). This transitional morphology suggests CPAP 3054 is not a hadrosaurid, but is nevertheless closely related to Hadrosauridae, having diverged shortly before the origin of this family. To properly address the relationships of CPAP 3054, we carried out the most comprehensive phylogenetic analysis of duck-billed dinosaurs to date, using a data matrix from a recent study (4) which we modified to correct scorings and include more characters, as well as *Gobihadros* and *Huehuecanauthlus,* which are important taxa with a morphology that is transitional to Hadrosauridae (42, 54), like CPAP 3054 (Methods). The South American duckbill *Lapampasaurus* was excluded because it is too fragmentary to confidently establish its affinities (adding this taxon also had no effect; see Methods). Our results for maximum parsimony analysis were unprecedentedly well-resolved, obtaining only 4 most parsimonious trees (1312 steps, CI: 0.396; RI: 0.821); further, all differences among these trees were among early-branching taxa at a node distant from CPAP 3054 (Fig. S6 shows the strict consensus of all 4 trees). Parsimony places CPAP 3054 outside of Hadrosauridae, but among transitional duckbills that are closest to this family (Fig. 5 and S6-S8; see Methods for selection and time calibration of this tree). Bayesian analysis also places CPAP 3054 as a close relative of Hadrosauridae, although its relations to other transitional duckbills were less resolved (Figs. S7 and S8). Resolution improved in the undated Bayesian analysis, which coincides with parsimony in that CPAP 3054 and the North American taxa *Lophorhothon* and *Huehuecanhautlus* are closer to Hadrosauridae than to other transitional duckbills (Fig. S7). Thus, these 3 taxa are likely the most relevant for understanding the origins of Hadrosauridae. However, both *Lophorhothon* and *Huehuecanhautlus* are represented by very partial materials (54, 61); currently, provides the best information, which is also bound to increase upon future excavations at the monotypic bonebed.

**Fig. 5.**
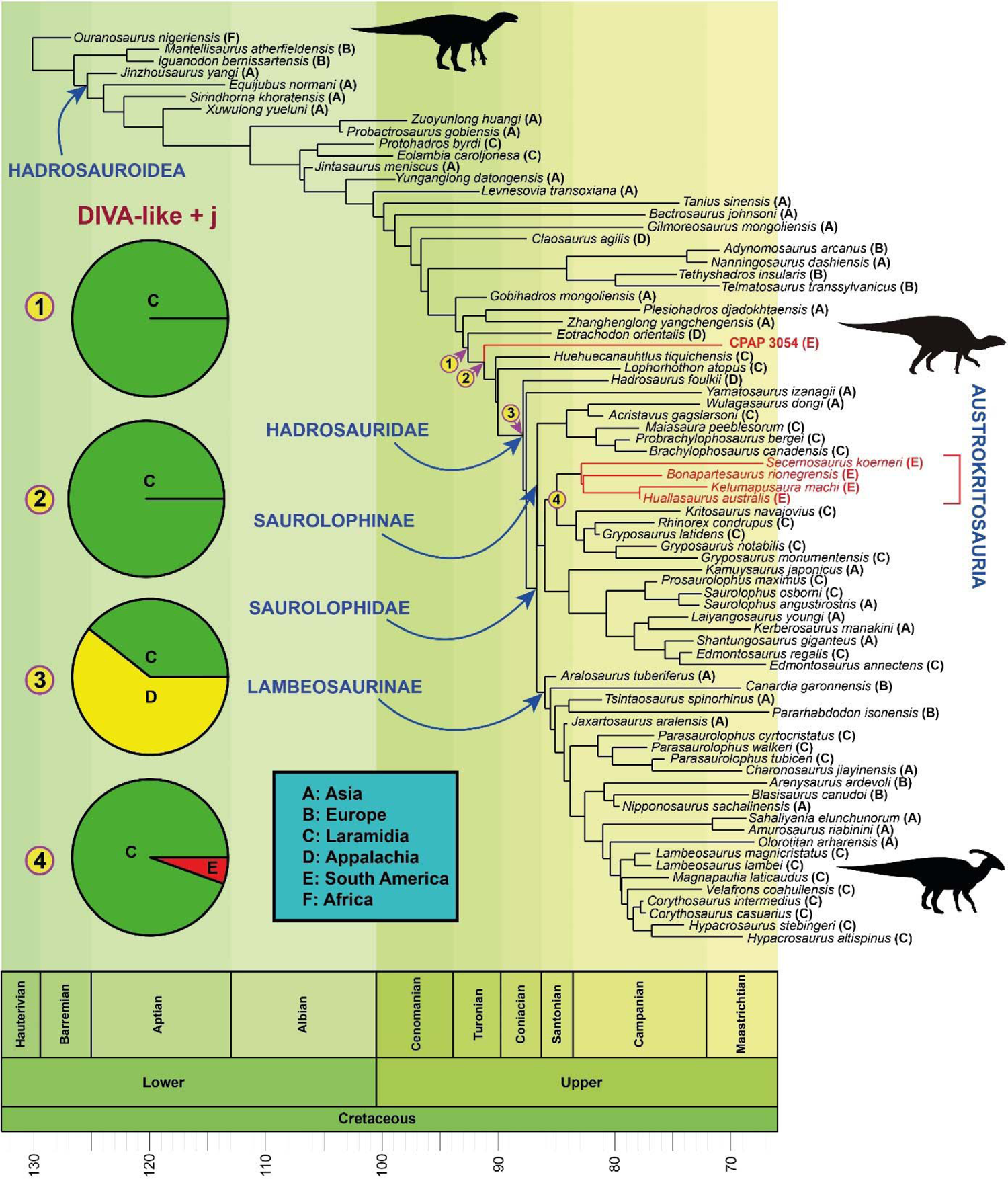
Phylogenetic relationships. Parsimony and Bayesian analyses recovered CPAP 3054 at a position transitional to Hadrosauridae, diverging shortly before the origin of this family; results of Parsimony are shown. The DIVA-like + j model (BioGeoBEARS) recovers Laramidia as the ancestral area for the most recent common ancestor of *Eotrachodon* and Hadrosauridae (node 1), as well as for the most recent common ancestor of CPAP 3054 and Hadrosauridae (node 2). Appalachia or Laramidia are recovered as ancestral areas for Hadrosauridae (node 3), and Laramidia as the ancestral area for Kritosaurini +Austrokritosauria (node 4). South American duck-billed dinosaurs are in red.

CPAP 3054 is not specially related to other South American duckbills, which are advanced forms of the family Hadrosauridae. Recent work has proposed that all other South American duckbills are saurolophine hadrosaurids, forming a monophyletic group that is the sister to the Kritosaurini, a tribe that inhabited Laramidia in North America (4). The monophyly of these South American hadrosaurids suggests that all members of this group descend from a single species that dispersed into South America. In our own analysis, despite inclusion of CPAP 3054 and some re-coding of South American duckbills, we recovered the same monophyletic group of South American saurolophine hadrosaurids in both Parsimony and Bayesian analyses, which was also recovered as sister to Kritosaurini by parsimony (Figs. 5, S6-S8). Although these South American hadrosaurids are related to Kritosaurini, they cannot be referred to this tribe: both parsimony and bayesian analyses do not support a nested position within the North American forms that are used to define Kritosaurini (62); thus, the current definition of Kritosaurini excludes these South American relatives. Given the need for a different and succinct term for the monophyletic group of “South American saurolophine hadrosaurids closely related to Kritosaurini”, we propose the name Austrokritosauria for the most inclusive clade that contains *Huallasaurus, Secernosaurus, Bonapartesaurus* and *Kelumapusaura,* but not *Gryposaurus*. Importantly, CPAP 3054 does not belong to this clade: placing CPAP 3054 as sister of Austrokritosauria requires a steep 15 extra steps in the parsimony analysis (Table S3). Therefore, CPAP 3054 represents a completely different lineage of duck-billed dinosaurs that independently colonized South America. As the most informative duck-billed dinosaur from far southern regions, CPAP 3054 also prescribes a reinterpretation of the record of partial fossils found in Southern Patagonia and Antarctica. This actually suggests hadrosaurids may have never arrived to these high austral latitudes. Partial remains in these regions can no longer be assumed a priori to belong to hadrosaurids like those of central and northern Patagonia: in fact, none of these remains show characters that are exclusive to Hadrosauridae, especially when we take into account the hadrosaurid-like features of CPAP 3054. In particular, two incomplete caudal centra from deposits of the Chorrillos Formation (Santa Cruz Province, Argentina) have been attributed to Hadrosauridae based on the hexagonal contour of their articular surfaces (11), but this trait is also present in CPAP 3054 (Fig. 4). An isolated and incomplete tooth from the Maastrichtian of Vega Island in the Antarctic Peninsula was attributed to Hadrosauridae, and tentatively to Hadrosaurinae (=Saurolophinae) because of its relatively symmetrical crown, with a single and nearly straight medial keel, and poorly developed denticles (13). Teeth are currently unknown for CPAP 3054 and most non-hadrosaurid duckbills, but the above characters were likely ancestral for Hadrosauridae, and may have been present in taxa close to this family; indeed, they are present in some maxillary teeth of *Eotrachodon* (44) and were likely present from before in evolution, since they are also observed in maxillary teeth of *Eolambia,* a much earlier non-hadrosaurid from North America (39). Also from the Antarctic peninsula, a metatarsal fragment found in Maastrichtian rocks of Seymour Island was attributed to Hadrosauridae (12), but its morphology has already been pointed out to be shared by other ornithopods, including those that inhabited these austral regions such as the Elasmaria (63) and now, non-hadrosaurid duckbills. In synthesis, southwards from central Patagonia (Chubut, 46°S), no remains can be reliably attributed to Hadrosauridae. This may add to other faunistic differences that have been proposed when comparing the fossil record of southern Patagonia and northern Patagonia (64), which could also relate to the presence of an archipelago at the time in southern South America, as a result of the marine transgressions that produced inland seas such as the Kawas Sea (14,15,16,17).

CPAP 3054 is the first non-hadrosaurid ever found in Gondwana, and its presence so far south poses a challenging biogeographic enigma. Any explanation implies remarkably long routes, large gaps in between with no records of non-hadrosaurid duckbills, and marine barriers that stopped most Laurasian dinosaurs; most likely, these marine barriers would have been breached by chains of islands rather than continuous land bridges (65), such that any dispersal of terrestrial animals must have involved swimming or “rafting” on debris (Fig. 6). The first American biotic exchange (between North America and South America) is proposed to have occurred during the latest Cretaceous (Campanian-Maastrichtian) (66). Mammals of North American origin were already diverse in the earliest Paleocene of South America, suggesting they must have crossed over earlier (67, 68). However, actual fossil evidence of exchanges during the latest Cretaceous is only provided by dinosaurs: the Austrokritosauria and the patagonian nodosaurid *Patagopelta* had North American ancestors (4, 69), while large titanosaurs in North America likely had ancestors in South America (70, 71).

**Fig. 6.**
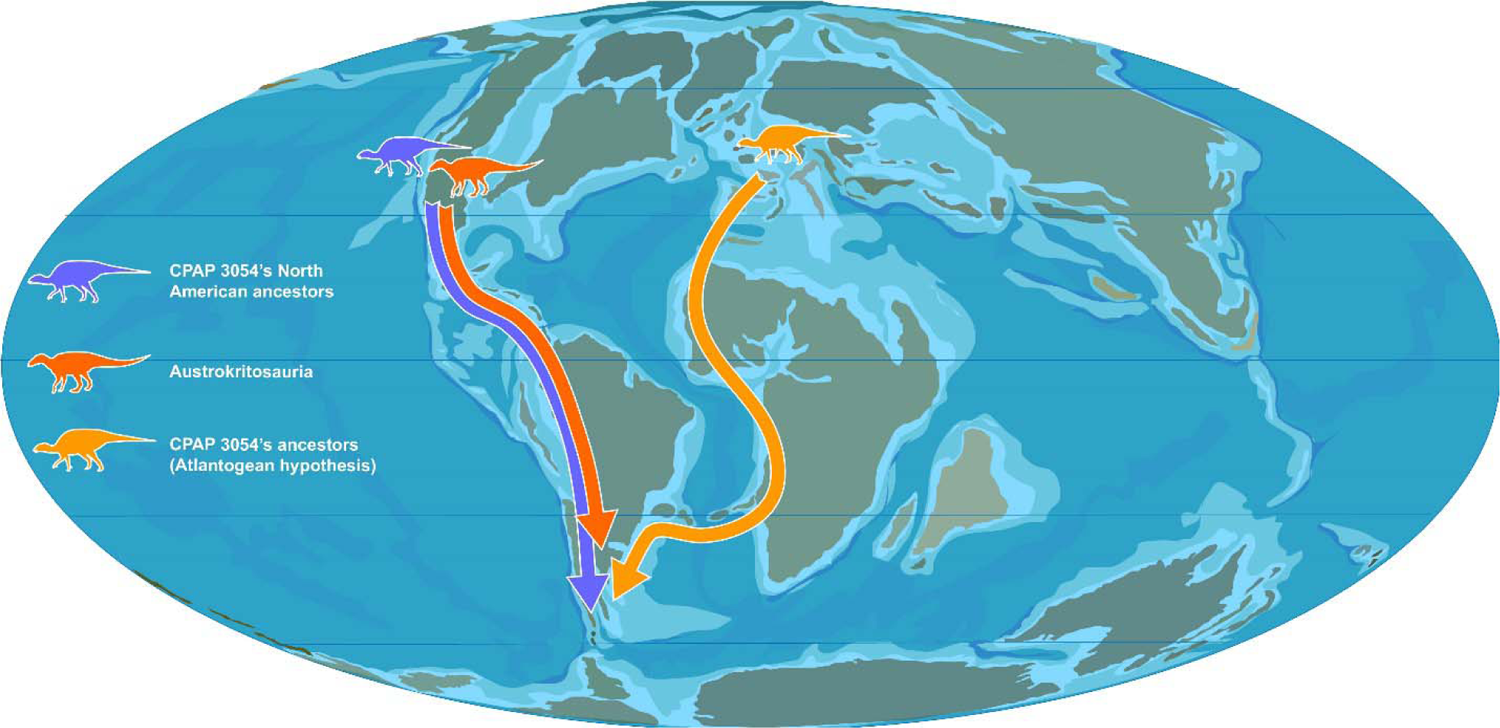
Biogeographic history of South American Duck-bill dinosaurs. Biogeographic statistics suggest that CPAP 3054’s ancestors arrived from North America (in purple). The alternative hypothesis, that CPAP 3054*’*s ancestors could have dispersed from Europe into South America via Africa (Atlantogean hypothesis, in yellow) was not supported by biogeographic statistics. Hadrosaurids related to the tribe Kritosaurini (in orange) would also have dispersed from North America, giving rise to the clade Austrokritosauria in South America. These hadrosaurids may have never reached the same high latitudes as CPAP 3054’s ancestors, possibly because they arrived later into South America. Paleogeographic drawing is based on information from the PALEOMAP project of Christopher Scotese and (10).

Another possibility is that non-hadrosaurids from Europe made it into Africa, and from there into South America (the “Atlantogean” route, 71; See Fig.5). This is a longer route that also implies crossing two marine barriers (rather than one), but exchanges through this route have been argued to be better supported than those between the Americas, based on numerous shared biotic components between Europe and South America during the latest Cretaceous (71). Whichever the case, duck-billed dinosaurs had the highest vagility (dispersal capacity) documented for any dinosaurs, with the greatest number of dispersal events that likely crossed marine barriers, namely: between Asia and Laramidia (9, 70); Appalachia and Laramidia (twice) (32, 72); Europe and Appalachia (50); Asia and Europe (twice) (50, 73); Europe and North Africa (5); Laramidia and South America (4); and South America and Antarctica (63). Duck-billed dinosaurs are also most often preserved near or within coastal environments (31; also the case for CPAP 3054; see Methods) and have been suggested to be apt swimmers, or even semi-aquatic (5, 51).

To formally test hypotheses on the biogeographical history of CPAP 3054, we reconstructed likely ancestral areas for all internal nodes of the phylogeny obtained from parsimony analysis, using BioGeoBEARS (74) as well as statistical Dispersal-Vicariance Analysis (s-DIVA) (75,76,77,78). The best-fit model in BioGeoBEARS was the DIVALIKE + j model (AICc: 182.21), which supports a strong role for events with founder effects (5, Table S4). This model provides clear-cut support for a Laramidian origin for the last common ancestor shared by CPAP 3054 and Hadrosauridae, as well as for the last ancestor shared with *Eotrachodon,* at the previous node (Figs. 5 and S12). The results of s-DIVA also support a North American origin, despite some differences, and decreased resolution: the last common ancestor shared by CPAP 3054 with Hadrosauridae is inferred in broad areas that combine Laramidia and/or Appalachia with South America, and the previous last ancestor shared with *Eotrachodon* is inferred as Appalachian (Fig. S13). As for Austrokritosauria, the results of both s-DIVA and BioGeoBEARS support a Laramidian origin. This is the first time that formal biogeographical statistical analysis is shown to support the occurrence during the Cretaceous of the first American biotic exchange, for CPAP 3054 and Austrokritosauria; similar studies may further confirm dispersal of nodosaurid ankylosaurs into South America, and of titanosaurian sauropods into North America.

Although Biogeographic statistics do not support an Atlantogean route (from Europe via Africa) for the ancestors of CPAP 3054, this could be a result of a sampling bias in Late Cretaceous faunas, which are very poorly known in Africa (5,79,80). Non-hadrosaurids were present in Europe in the Late Cretaceous (35, 50), when lambeosaurine hadrosaurids managed to disperse from Europe into Africa (5), which suggests non-hadrosaurids could have done the same. To explore this, we added a hypothetical African taxon of a non-hadrosaurid duckbill to our BioGeoBEARS and s-DIVA analyses, placing it between CPAP 3054 and *Eotrachodon*, or as sister to CPAP 3054. A North American origin continued to be best supported, suggesting that this conclusion will not be easily overturned, even if such discoveries are made in Africa. (Figures S12-S15, Tables S5 and S6, see Methods and Supplementary Material for details).

Some dinosaur clades that are present in both North and South America have been shown to reflect ancient cosmopolitan origins, before the final separation of Gondwana and Laurasia (81,82,83,84,85,86,87). Although CPAP 3054 is not a hadrosaurid, such a scenario would require for a much earlier divergence near the origin of duck-billed dinosaurs (Hadrosauroidea) in the late Hauterivian (130 mya), when biotic exchanges between Europe and Gondwana were still possible (2012). Such a basalmost position for CPAP 3054 would require a steep 45 extra steps (Table S3). Given the actual phylogenetic position of CPAP 3054, to insist on ancient cosmopolitan roots would imply that duckbills close to Hadrosauridae had already originated in the Hauterivian, despite a gap of about 45 million years in their worldwide fossil record (Fig. 5).

We conclude that CPAP 3054 is very likely descended from North American non-hadrosaurids that are transitional to Hadrosauridae, of the likes of *Eotrachodon, Lophorhothon* and *Huehuecanhautlus*. These were also the last non-hadrosaurids of North America, where they were replaced by hadrosaurids: *Eotrachodon* and *Huehuecanhautlus* are of Santonian age; only *Lophorothon* may have lived in the Campanian, given its large chronostratigraphic uncertainty, ranging from the latest Santonian (83,6 mya) to the late Campanian (75,8 mya) (32,54,61). Non-hadrosaurids are absent in the abundant and well-studied record of North American dinosaurs of the latest Campanian and Maastrichtian, strongly suggesting they had become locally extinct (41, 88). However, at some point before that, the non-hadrosaurid ancestors of CPAP 3054 managed to leave North America, surviving into the Maastrichtian as distant relict populations in Subantarctic Chile.

Given that CPAP 3054 is found so far south, and that no hadrosaurid remains can be confirmed from Subantarctic and Antarctic regions, this pattern suggests the ancestors of CPAP 3054 arrived earlier than those of Austrokritosauria, giving the former a head-start in reaching more austral regions. Hadrosaurids may have not had enough time to reach this far south before the K-Pg mass extinction. This interpretation is allowed by the divergence times in our time-calibrated tree: CPAP 3054 would have last shared an ancestor with North american forms around 91 mya (Turonian), while the Austrokritosauria would have shared their last ancestor with the North American Kritosaurini around 85 mya (Santonian; see Methods for calibration details).

Another reason to suspect that hadrosaurids may have never arrived into Subantarctic and Antarctic regions is that they tended to replace non-hadrosaurids in those regions where they co-existed. The niches of hadrosaurids likely overlapped onto those of non-hadrosaurids, while non-hadrosaurids like CPAP 3054 were often smaller-sized, and had smaller tooth plates with fewer tooth positions, taking up a smaller proportion of the jaw (39,40,47,48). Given the importance of the tooth-jaw apparatus, this could explain why in the Maastrichtian non-hadrosaurids had disappeared in North America (41, 88), while a single species is known from Asia (89, 90). Non-hadrosaurids like *Telmatosaurus* persisted into the Maastrichtian of Europe, but this is argued to have resulted from their geographic isolation in islands (35, 50). This is also a feasible explanation for CPAP 3054, given marine transgression events at the time in southern South America.

Although the radiation of the Hadrosauridae in North America and Central China coincides with a decline in dinosaur biodiversity previous to the K-Pg asteroid impact, the presence of Hadrosauridae cannot be confirmed in southern Patagonia and Antarctica, while New Zealand and Australia may have had no duckbills at all (hadrosaurid or not), with exclusively endemic gondwanan faunas right until the K-Pg mass extinction (91). It remains to be seen if declines in dinosaur biodiversity were a global phenomenon; if so, they did not correlate everywhere with the radiation of the Hadrosauridae. Alternatively, if declines in dinosaur biodiversity did not occur in austral regions, their ecosystems were nevertheless equally vulnerable to mass extinction upon the asteroid impact, and would suggest a greater role of the impact itself as a kill mechanism (92).

## Materials and Methods

### Experimental model and subject details

The anatomical information used for both our comparisons and phylogenetic analyses were obtained from the literature and from direct observation of specimens housed in public repositories. The specimens mentioned below were observed directly:

– *Huallasaurus australis*, Natural History Museum Bernardino Rivadavia, Buenos Aires, Argentina (MACN-RN 02, MACN-RN 142, MACN-RN 142B, MACN-RN 143, MACN-RN 144, MACN-RN 145, MACN-RN 146, MACN-RN 826).
– *Secernosaurus koerneri*, FMNH, The Field Museum, Chicago, USA (FMNH PP13423).
– *Bonapartesaurus rionegrensis*, Vertebrate Paleontology Collection at the National University of Tucuman and the Fundación Miguel Lillo (Tucuman, Argentina), (MPCA-Pv SM2).

### Specimen provenance and geological Setting

The specimens studied here were found in the Río de las Chinas Valley located 350 km to the north of the city of Punta Arenas, Ultima Esperanza Province (Magallanes Region, Chile). The studied outcrops extend along the valley in N-S direction, near to the international border with Argentina (Fig.1). The fossil-bearing levels are assigned to the Dorotea Formation (Upper Campanian to Danian; 29,93,94), which corresponds to the upper section of the continental-marine sedimentary succession that fills the Magallanes/Austral Basin. This basin was formed during the break-up of Gondwana and the opening of the Atlantic Ocean (95,96,97). This succession conformably overlies the Campanian to lower Maastrichtian Tres Pasos Formation (98, 99) and is unconformably overlain by the Paleogene sequences of the Man Aike/Río Turbio formations of middle to late Eocene age (100, 101). The Dorotea Formation represents a transition between shallow marine and continental environments (102,103,104,105,106,107,108,109,110,111,112). Specifically, it has been interpreted as a transition from a shallow marine shelf-edge to tide-dominated delta and fluvial systems (106,110,111,112,113,114).

The Dorotea Formation mainly comprises greenish-gray and reddish-brown sandstones with abundant conglomerate and siltstone lenses, thin beds of sandy calcareous concretions and mudstones of 900 to 1200 m thickness (98,110,115). Along the succession there are fossil-bearing levels with bivalves, ammonites, gastropods, sharks, plesiosaurs, mosasaurs, frogs, turtles, dinosaurs, mammals, fossil wood and leaves (30,87,98,110,111,116,117,118,119,120,121,122).

The material here studied comes from the upper section of the Dorotea Formation in the Río de las Chinas Valley, consisting of brown sandy mudstone (60 cm thickness) with coal lenses and reddish-brown fine-grained sandstone (20 cm thickness) levels (CHA-2, Fig. 1). Numerous small to medium-sized hadrosauroid bones were discovered in the sandy mudstone level and larger bones in the sandstones. The bones are preserved three-dimensionally and show linear and angular fragmentation. In Addition, a 0.5 m thick silicified trunk was discovered in the surface debris on a topographic slope. Apparently, this layer with hadrosaur bones extends laterally for about 5 km to the northeast, exposing itself at several points in the valley. The most notable differences have to do with the grain size at the different points where this layer is exposed. Further stratigraphic studies are needed to determine whether this entire layer originated from a single depositional event, and whether bones exposed at other points also correspond to CPAP 3054. The depositional environment of these levels with hadrosauroid bones are interpreted as the distal section of a floodplain in a continental environment (fluvial channels), with lower energy. An age estimation using U-Pb detrital zircon for the levels above the hadrosauroid horizons provided values of 71.7 ± 1.2 Mya and 70.5 ± 4.5 Mya (29, 30), supporting an early Maastrichtian age (Late Cretaceous) for the fossil-bearing levels.

### Extraction and technical preparation

After the initial finding of fossil bones in situ at the exposed surface, we identified the bearing-bone layer. The level was excavated in detail and several more specimens appeared randomly distributed. Bone remains were extracted individually or as a set of bones, using plaster jackets for protection and transport. Some bones were prepared in the Paleobiology Laboratory of the Chilean Antarctic Institute (INACH), located in Punta Arenas, Magallanes Region, while most of them were prepared in the Paleontology Laboratory of the Faculty of Sciences of the University of Chile, located in Santiago de Chile. The bones were prepared using air scribes, helped with tips and needles, in some cases under a binocular microscope. Glues such as paraloid (B-72) and cyanoacrylate were used to stabilize the samples. Finally, each element was numbered and conditioned for their preservation in the collection of INACH with the acronym CPAP.

### Character Dataset and taxa

For the phylogenetic analyses (and subsequent biogeographical analyses), we used a modified version of the matrix of a recent phylogenetic analysis of all duck-billed dinosaurs (Hadrosauroidea) by (4), with the most extensive character and taxon sample published to date. We rescored/redefined 20-character states:

1. *Secernosaurus koerneri* (character 258). Changed from 1 to ?, since the distal half of the scapula is not preserved.
2. *Secernosaurus koerneri* (character 259). Changed from 0 to ?, since the scapula is incomplete.
3. *Secernosaurus koerneri* (character 278). Changed fromto 1, based on fig. 12 of (123).
4. *Secernosaurus koerneri* (character 290). Changed from 0 to 1, based on calculated proportions.
5. *Secernosaurus koerneri* (character 293). Changed from 2 to 1 since the supraacetabular process does not extend far enough ventrally to reach the middle of the iliac blade.
6. *Secernosaurus koerneri* (character 303). Changed from 0 to 1, as the thickening of the postacetabular process is due to its dorsomedial rotation.
7. *Secernosaurus koerneri* (character 312). Changed from 1 to 0, since in our opinion, the pubic peduncle of the pubis is shorter and more robust compared with other hadrosauroids, see (47).
8. *Bonapartesaurus rionegrensis* (character 247). Changed fromto 0, since the chevrons are shorter than the caudal neural spines.
9. *Bonapartesaurus rionegrensis* (character 291). Changed from 0 to 1, since the apex of the supraacetabular process is in an anterodorsal position with respect to the caudal tuberosity of the ischial peduncle of the ilium.
10. *Bonapartesaurus rionegrensis* (character 335). Changed from 1 to 0, based on the original description given in (9).
11. *Kelumapusaura machi* (character 291). Changed from 0 to 1 since the apex of the supraacetabular process is in an anterodorsal position with respect to the caudal tuberosity of the ischiadic peduncle of the ilium.
12. *Huallasaurus australis* (character 287). Changed from 0 to 1 based on our observations.
13. CPAP 3054 (character 263). Redefinition (new character state added). A fourth state (3) was added for the pseudoacromion process, which refers to the ventral curvature of this structure.
14. *Laiyangosaurus youngi* (character 17). Changed from 2 (undefined character in character list) to 1. Based on the description of (124).
15. *Parasaurolophus walkeri* (character 89). Changed from 2 (undefined character in character list) to 1 based on (125).
16. *Olorotitan arharensis* (character 89). Changed from 2 (undefined character in character list) to 1 based on (125).
17. *Hypacrosaurus altispinus* (character 89). Changed from 2 (undefined character in character list) to 1 based on (125).
18. *Lambeosaurus lambei* (character 91). Changed from 2 (undefined character in character list) to 1 based on (125).
19. *Lambeosaurus magnicristatus* (character 91). Changed from 2 (undefined character in character list) to 1 based on (125).
20. *Hypacrosaurus altispinus* (character 91). Changed from 2 (undefined character in character list) to 1 based on (125).

In addition, we added 5 new characters and asides from CPAP 3054:

Character 361 – Maxilla, ectopterygoid ridge: continuous with jugal tubercle (0) or discontinuous (1). Added from (5), Character 244.

Character 362 – Maxilla, ectopterygoid ridge: ridge extends to ventral jugal tubercle (0) main part of shelf lies distinctly below it (1). Added from (5), Character 245.

Character 363 – Maxilla, neurovascular foramina: a single very large foramen below the jugal, with a second, smaller foramen below it: absent (0); present (1). Added from (5), Character 247.

Character 364 – Maxilla, posterior dentigerous process: dentigerous margin straight (0); posterior downturn (1) strong posterior downturn, i.e., “boomerang” shape (2). Added from (5), Character 340.

Character 365 – Humerus, shape of the deltopectoral crest apex: well rounded (0); extending abruptly to produce a prominent angular profile (1). Added from (126), Character 37; following (40), Character 221.

We also added *Gobihadros mongoliensis* and *Huehuecanauhtlus tiquichensis*. As in previous work (4), *Lapampasaurus* was left out from the results because its extremely partial remains (only 11 of 365 characters) preclude any significant inference of its phylogenetic affinities. Inclusion in our analysis did not alter the topologies of the 4 MPT’s and placed this south american taxon within Saurolophinae, but retrieved it closest to the japanese *Kamuysaurus*.

The character distribution was modified with Mesquite 2.75 (127). The characters were coded based on bones belonging to adult or subadult individuals of CPAP 3054. The character states of juvenile individuals were only coded for characters that do not show significant ontogenetic variation. We modified the coding of some characters of *Huallasaurus australis* based on direct observation by two of the co-authors (SSA and PC-C). We also modified the scoring of some characters of *Secernosaurus koerneri* and *Bonapartesaurus rionegrensis* based on published images and direct observation by AOV (*Secernosaurus koerneri*) and PC-C (*Bonapartesaurus rionegrensis*). The resulting matrix included 77 species-level taxonomic units (four non-hadrosauroid iguanodontians and 73 hadrosauroids) scored for 365 equally weighted characters.

### Parsimony analysis

The maximum parsimony analysis was carried out in TNT version 1.5 (128). The non-hadrosauroid iguanodontian *Ouranosaurus nigeriensis* was selected as the outgroup. A heuristic search of 1000 replicates of Wagner trees using random additional sequences was performed, followed by branch swapping by tree-bisection-reconnection, retaining 100 trees per replicate. All characters were treated as equally weighted and unordered. For all the trees, Bremer support (129) was calculated for each node to assess its robustness by computing decay indices in TNT using the script Bremer.run (130). Bootstrap proportions (131) were also calculated with TNT, setting the analysis for 5000 replicates using heuristic searches (see Fig. S6).

### Templeton tests

We used a Templeton test to assess the significance of different hypotheses where CPAP 3054 was forced into alternative phylogenetic positions, evaluating changes in the number of steps. We randomly used MPTs obtained from the unconstrained analysis to compare with the forced topologies, applying a one-sided Wilcoxon signed-rank test to the differences in character transformations between trees. We assessed the forced positions of CPAP 3054 as the basalmost hadrosauroid (1), and as a member of the Austrokritosaurini (2). (Table S3) We assessed whether the difference in steps between the original and forced hypotheses was significant using Templeton Tests (132), implemented with a script in TNT 1.5 (133).

### Selection and Time calibration of trees obtained from Parsimony analysis

Only 4 trees were obtained from parsimony analysis with the standard option to collapse zero-length branches. Close relationships of CPAP 3054 were fully resolved: differences among trees were all in taxa distant from CPAP 3054, represented in a single polytomy at a basal node (see Fig. S6). We selected one of the 4 trees to show the relationships of CPAP 3054, and for biogeographic analyses with BioGeoBEARS, which require a fully resolved tree (see below). We picked the only tree with no zero-length branches (no polytomies), which is shown in Fig. 5 (tree number 0 in TNT). To time-calibrate this tree, we ran a tip-dated Bayesian analysis with the entire topology constrained to this tree; we kept the same priors and MCMC parameters than in the unconstrained Bayesian Analysis (see below) and used the resulting maximum clade compatibility tree (MCCT). To obtain the MCCT tree we used the posterior files of MrBayes and the obtainDatedPosteriorTreesMrB() function of the R package paleotree v3.4.4 (134). Table below shows FADs (First Appearance Data) and LADs (Last Appearance Data), which were used for the time calibration of the phylogenetic tree according to chronostratigraphic uncertainty ranges (oldest possible age and youngest possible age).

FADs (First Appearance Data) and LADs (Last Appearance Data):

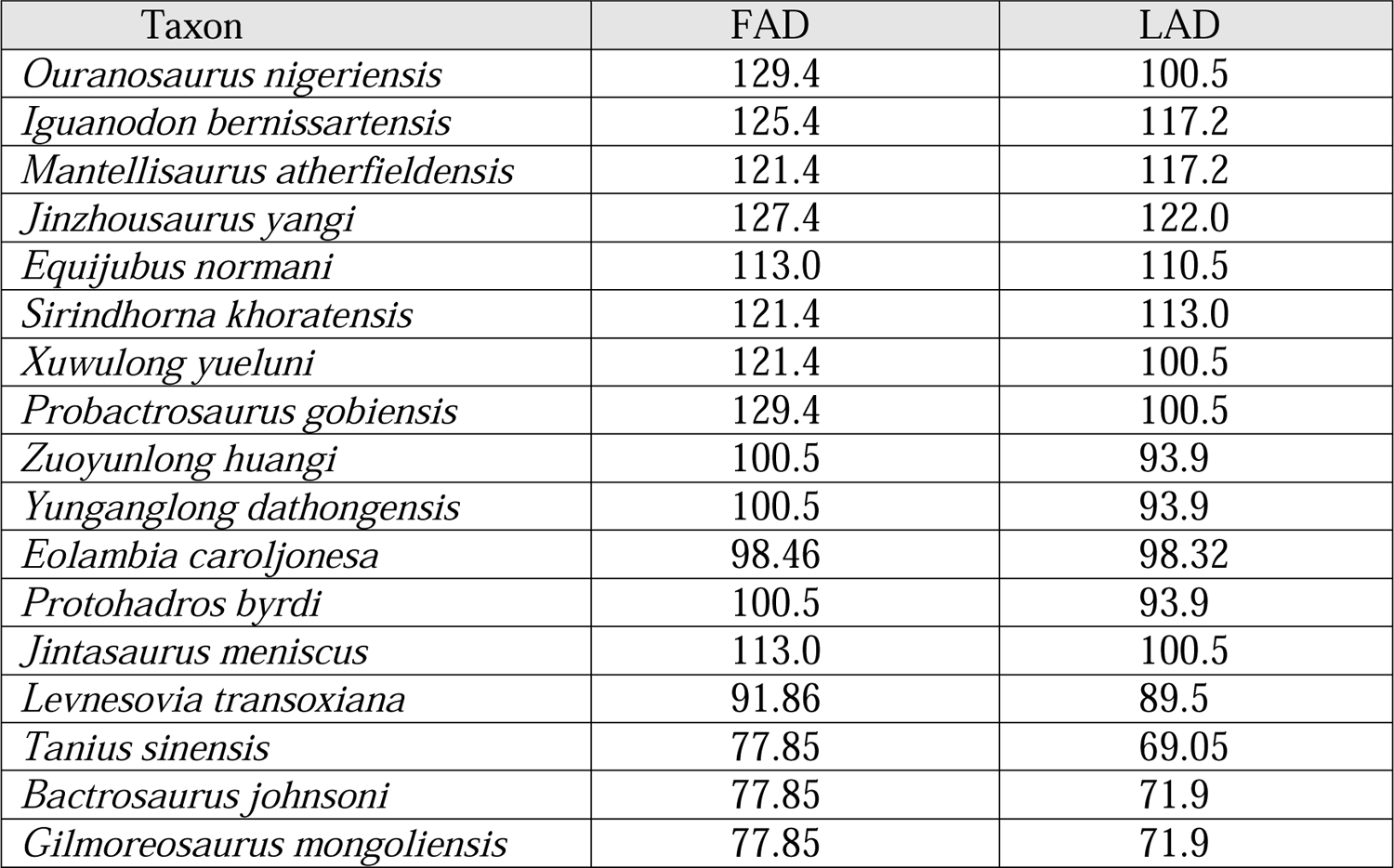
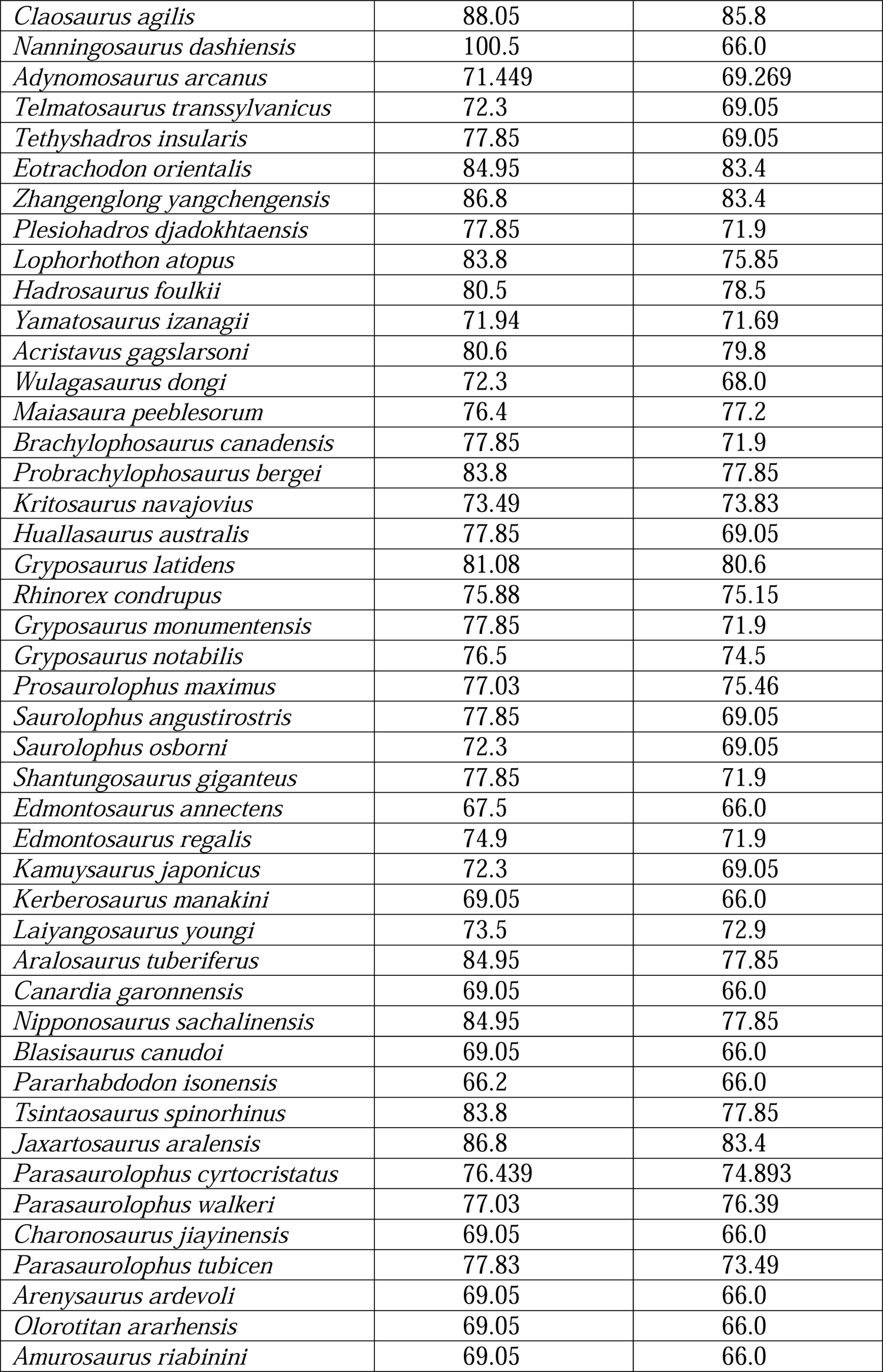
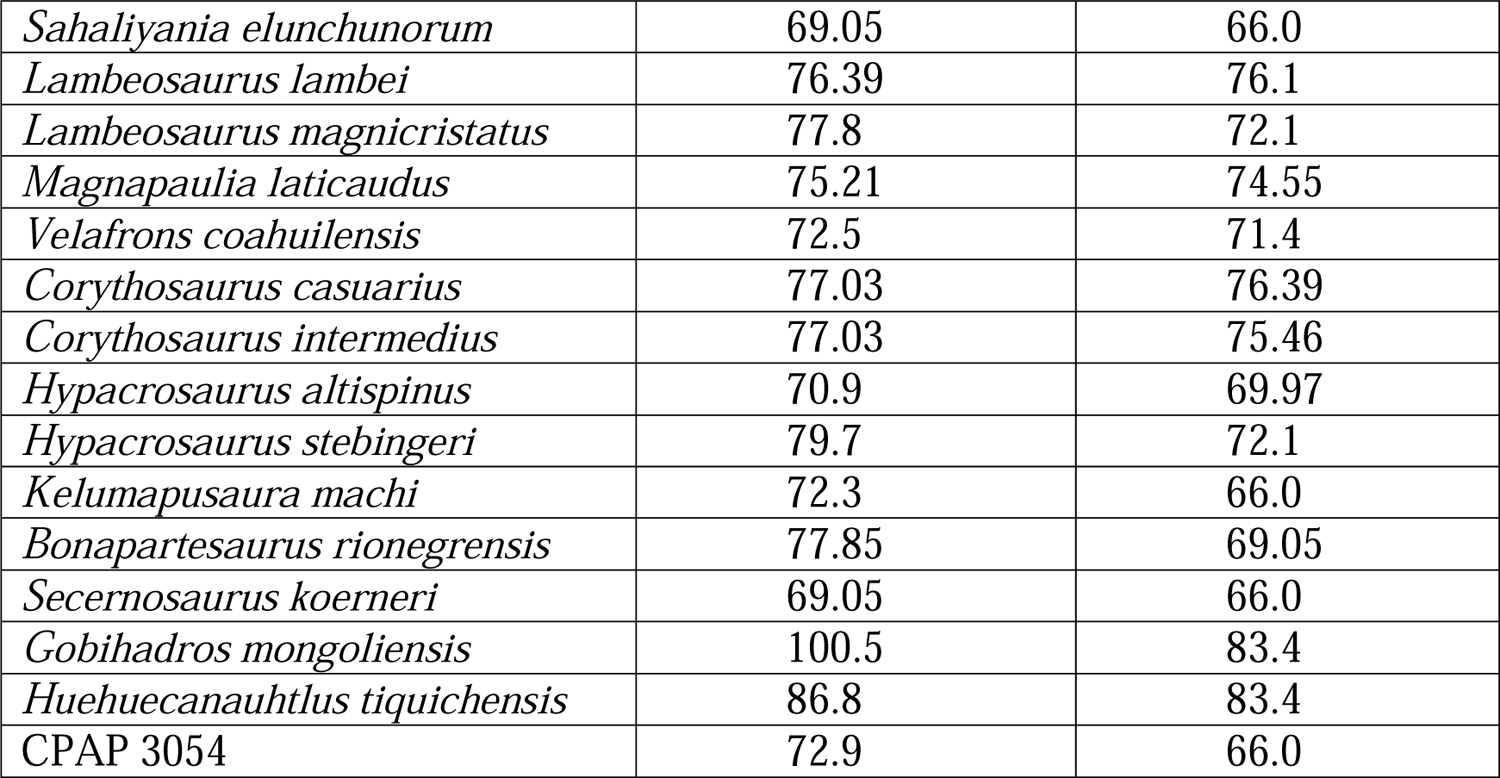

| Taxon | FAD | LAD |
| --- | --- | --- |
| *Ouranosaurus nigeriensis* | 129.4 | 100.5 |
| *Iguanodon bernissartensis* | 125.4 | 117.2 |
| *Mantellisaurus atherfieldensis* | 121.4 | 117.2 |
| *Jinzhousaurus yangi* | 127.4 | 122.0 |
| *Equijubus normani* | 113.0 | 110.5 |
| *Sirindhorna khoratensis* | 121.4 | 113.0 |
| *Xuwulong yueluni* | 121.4 | 100.5 |
| *Probactrosaurus gobiensis* | 129.4 | 100.5 |
| *Zuoyunlong huangi* | 100.5 | 93.9 |
| *Yunganglong dathongensis* | 100.5 | 93.9 |
| *Eolambia caroljonesa* | 98.46 | 98.32 |
| *Protohadros byrdi* | 100.5 | 93.9 |
| *Jintasaurus meniscus* | 113.0 | 100.5 |
| *Levnesovia transoxiana* | 91.86 | 89.5 |
| *Tanius sinensis* | 77.85 | 69.05 |
| *Bactrosaurus johnsoni* | 77.85 | 71.9 |
| *Gilmoresaurus mongoliensis* | 77.85 | 71.9 |
| *Claosaurus agilis* | 88.05 | 85.8 |
| *Nanningosaurus dashiensis* | 100.5 | 66.0 |
| *Adynomosaurus arcanus* | 71.449 | 69.269 |
| *Telmatosaurus transsylvanicus* | 72.3 | 69.05 |
| *Tethyshadros insularis* | 77.85 | 69.05 |
| *Eotrachodon orientalis* | 84.95 | 83.4 |
| *Zhangenglong yangchengensis* | 86.8 | 83.4 |
| *Plesiohadros djadokhtaensis* | 77.85 | 71.9 |
| *Lophorhodon atopus* | 83.8 | 75.85 |
| *Hadrosaurus foulkii* | 80.5 | 78.5 |
| *Yamatosaurus izanagii* | 71.94 | 71.69 |
| *Acristavus gagslarsoni* | 80.6 | 79.8 |
| *Wulagasaurus dongi* | 72.3 | 68.0 |
| *Maiasaura peeblesorum* | 76.4 | 77.2 |
| *Brachylophosaurus canadensis* | 77.85 | 71.9 |
| *Probrachylophosaurus bergei* | 83.8 | 77.85 |
| *Kritosaurus navajovius* | 73.49 | 73.83 |
| *Huallasaurus australis* | 77.85 | 69.05 |
| *Gryposaurus latidens* | 81.08 | 80.6 |
| *Rhinorex condrupus* | 75.88 | 75.15 |
| *Gryposaurus monumentensis* | 77.85 | 71.9 |
| *Gryposaurus notabilis* | 76.5 | 74.5 |
| *Prosaurolophus maximus* | 77.03 | 75.46 |
| *Saurolophus angustirostris* | 77.85 | 69.05 |
| *Saurolophus osborni* | 72.3 | 69.05 |
| *Shantungosaurus giganteus* | 77.85 | 71.9 |
| *Edmontosaurus annectens* | 67.5 | 66.0 |
| *Edmontosaurus regalis* | 74.9 | 71.9 |
| *Kamuysaurus japonicus* | 72.3 | 69.05 |
| *Kerberosaurus manakini* | 69.05 | 66.0 |
| *Laiyangosaurus youngi* | 73.5 | 72.9 |
| *Aralosaurus tuberiferus* | 84.95 | 77.85 |
| *Canardia garonnensis* | 69.05 | 66.0 |
| *Nipponosaurus sachalinensis* | 84.95 | 77.85 |
| *Blasisaurus canudoii* | 69.05 | 66.0 |
| *Pararhabdodon isonensis* | 66.2 | 66.0 |
| *Tsintaosaurus spinorhinus* | 83.8 | 77.85 |
| *Jaxartosaurus aralensis* | 86.8 | 83.4 |
| *Parasaurolophus cyrtocristatus* | 76.439 | 74.893 |
| *Parasaurolophus walkeri* | 77.03 | 76.39 |
| *Charonosaurus jiayinensis* | 69.05 | 66.0 |
| *Parasaurolophus tubicen* | 77.83 | 73.49 |
| *Arenysaurus ardevoli* | 69.05 | 66.0 |
| *Olorotitan ararhensis* | 69.05 | 66.0 |
| *Amurosaurus riabinini* | 69.05 | 66.0 |
| *Sahaliyana elunchunorum* | 69.05 | 66.0 |
| *Lambeosaurus lambei* | 76.39 | 76.1 |
| *Lambeosaurus magnicristatus* | 77.8 | 72.1 |
| *Magnapaulia laticaudus* | 75.21 | 74.55 |
| *Velafrons coahuilensis* | 72.5 | 71.4 |
| *Corythosaurus casuarius* | 77.03 | 76.39 |
| *Corythosaurus intermedius* | 77.03 | 75.46 |
| *Hypacrosaurus altispinus* | 70.9 | 69.97 |
| *Hypacrosaurus stebingeri* | 79.7 | 72.1 |
| *Kelumapusaura machi* | 72.3 | 66.0 |
| *Bonapartesaurus rionegrensis* | 77.85 | 69.05 |
| *Secernosaurus koernerii* | 69.05 | 66.0 |
| *Gobihadros mongoliensis* | 100.5 | 83.4 |
| *Huehuecanauhtlus tiquichensis* | 86.8 | 83.4 |
| CPAP 3054 | 72.9 | 66.0 |

### Bayesian phylogenetic analyses (undated and tip-dated)

We conducted an undated Bayesian analysis in MrBayes v3.27a (135). As in the parsimony analysis, the outgroup was the non-hadrosauroid iguanodontian *Ouranosaurus nigeriensis*. We used the Mkv + Γ model of morphological evolution with ascertainment bias correction (136), setting the coding as variable and with six rate categories for the gamma distribution. We used two independent runs for the analysis with 40 million Markov chain Monte Carlo (MCMC) generations, sampling every 1000 generations and discarding 50% of the samples as burn-in. The posterior distributions for each parameter were checked to confirm that the effective sample size was > 200 and the deviation of split frequencies was below 0.05.

In addition, we performed a tip-dated Bayesian analysis in MrBayes v3.27a (135) using the same settings for the model of morphological evolution (see above). The root of the tree was calibrated using an exponential distribution with a minimum age of 129.4 mya and a mean of 139.4 mya. The minimum age of the root was based on the maximum age of the non-hadrosauroid iguanodontian *Ouranosaurus nigeriensis*. For calibrating the tree tips, we used a uniform distribution based on the radiometric age uncertainty of the fossils scored in the morphological matrix (137)

We used a normal distributed clock rate prior, with a mean of 0.02253 and standard deviation of 0.00839. This prior was extracted from the previously obtained undated Bayesian consensus tree and by using the packages ape (138) and fitdistrplus (139) from R 2022.07.2+576 (140) following the methodology described in (141). The best model following the Bayesian information criterion was the gamma distributed clock rate (shape = 5.92176, rate = 262.84887). However, preliminary analyses of the data showed problems of convergence between the two independent runs using this model, therefore the normal distributed model was preferred for our analysis. We used the FBD model using an exponential net diversification prior with rate 1, a beta fossil sampling proportion prior with shape parameters α = 1 and β = 1, a beta turnover prior with shape parameters α = 1 and β = 1, and an extant sampling proportion of 1. We modeled branch rate variation using the Independent Gamma Rate (IGR) relaxed clock model with an exponential distribution of rate 10. We used diffuse priors for the FBD model and the clock variance, which reflect our prior uncertainty in the distribution of these parameters. We used two independent runs for the analysis with 40 million Markov chain Monte Carlo (MCMC) generations, sampling every 1000 generations and discarding 50% of the samples as burn-in. The posterior distributions for each parameter were checked to confirm that the effective sample size was > 200 and the deviation of split frequencies was below 0.05.

In the 50% majority-rule consensus tree of the tip-dated Bayesian analysis, CPAP 3054 is placed in a polytomy with *Eotrachodon*, *Zhangenlong* and *Plesiohadros* (Fig. S8). Resolution improved in the undated Bayesian analysis, when geological ages were not allowed to influence the topology; as in the parsimony analysis, CPAP 3054 was retrieved closer to the clade that includes *Lophorhothon*, *Huehuecanauthlus* and Hadrosauridae than to *Eotrachodon*, *Zhangenlong* or *Plesiohadros* (Fig. S7). This difference between tip-dated and undated analysis may result from the fact that the ages of the nearby *Nanningosaurus* and *Gobihadros* are very uncertain, with large possible temporal ranges. Regardless, in both approaches, many polytomies persisted regarding the close relationships of CPAP 3054. Even in the maximum posterior probability trees (not shown), most relationships were recovered in less than 50% of trees. We therefore decided not to select any tree resulting from Bayesian phylogenetic analysis for the purposes of biogeographic analyses (which require fully resolved trees).

### Statistical Biogeographic analyses

We implemented a statistical Dispersal-Vicariance Analysis (s-DIVA75,76,77,78) in RASP 4.2 (78) to reconstruct the ancestral areas for all internal nodes of the phylogeny obtained in our parsimony analysis. Six general areas were considered: Asia (A), Europe (B), Laramidia (C), Appalachia (D), South America (E) and Africa (F). Prior to the s-DIVA, we performed a new phylogenetic analysis in TNT 1.5 to obtain the phylogenetic trees that were used in this analysis. The phylogenetic analysis recovered 6 most parsimonious trees without collapsing zero-length branches (CI: 0.369, RI: 0.821), to obtain fully resolved trees. The 6 trees were loaded in RASP 4.2, as well as the strict consensus tree, which was used for the graphical representation of the s-DIVA results.

The “Allow Reconstruction (Slow)” option was selected to implement the method of calculating the frequency of ancestral states for each node (142). The Supplementary Material contains the details of the results.

We also used the R package BioGeoBEARS v.1.12 (74) as a second approach for estimating the biogeographic history of duck-billed dinosaurs and the ancestral areas for CPAP 3054. This method evaluates alternative models for extinction, dispersal, cladogenesis and founder effect (143). BioGeoBears requires fully resolved time-calibrated trees. We did not use any tree obtained from Bayesian inference since most relationships of CPAP 3054 remained unresolved in the 50% majority rule tree (144).

Therefore, we used the only tree with no zero-length branches obtained from parsimony analysis, which we time-calibrated by running a tip-dated Bayesian analysis with the entire topology constrained to this tree (see above). The terminal taxa were coded for the following biogeographical provinces: Asia, Europe, Laramidia, Appalachia, South America, and Africa. We used a dispersal matrix that considers different dispersal probabilities between land masses. The dispersal weights between geographical areas were obtained from (5). We evaluated six different models: DIVALIKE and DIVALIKE + j (75), DEC and DEC + j (145), and BAYAREALIKE and BAYAREALIKE + j (146).

We permitted only four simultaneous areas. These models were compared using their AICc (Corrected Akaike Information Criterion). Values of log-Likelihood (lnL), Dispersal (d), Extinction (e), Founder effect (j), Corrected Akaike Information Criterion (AICc), and AICc Weight (AICc wt) scores from each model implemented. The model with the lowest AICc value (and higher AICc + wt) was selected as the model with the best fit to the data.

## Acknowledgments

To Edwin González, Sebastián Jiménez, Juan Pablo Venegas, Roy Fernández, Juan Pablo Guevara, María Jesús Ortuya, Sebastían Garrido, Viviana Lobos, Roberto Yury, Alejandra Manríquez, Marcelo Isasi, Marcela Milani, Sebastián Apesteguía, Jared Amudeo, Valentina Poblete, Cynthia Cabezas, Antonia Atisha, Daniela Flores and Claudio Bravo for their valuable help in the excavations and/or technical preparation of the CPAP 3054 bones; to Estancia Cerro Guido and especially the Matetic, Simunovic and Reyes families for granting access and important logistical support in the field; to the Consejo de Monumentos Nacionales (National Monuments Council) of the Chilean Ministry of Culture, Arts and Heritage for fieldwork permits to Marcelo Leppe. We thank William F. Simpson (Field Museum of Natural History, USA) for sending us photographs of the scapula of the holotype of *Secernosaurus koerneri*. To Martín Ezcurra (Museo Argentino de Ciencias Naturales “Bernardino Rivadavia”, Buenos Aires, Argentina) for clarifying doubts about calibration of phylogenetic trees. Finally, we thank Agustín Martinelli (Museo Argentino de Ciencias Naturales “Bernardino Rivadavia”, Buenos Aires, Argentina) for allowing us access to the materials of *Huallasaurus australis*.

## Author contributions

Conceptualization: J.A.M., A.O.V., H.P., P.C.-C.

Methodology: J.A.M., A.O.V., H.P., S.S.-A., J.P.-L., L.M., V.M., C.S.-G., P.C.-C

Investigation: J.A.M., A.O.V., H.P., S.S.-A., L.M., J.P.P., D.B., V.M., C.S.-G., P.C.-C.

Resources: A.O.V., M.L., P.C.-C.

Data Curation: J.A.M., H.P., J.P.-L.

Formal Analysis: J.A.M., H.P., J.P.-L.

Visualization: J.A.M., A.O.V., H.P., L.M., V.M.

Writing—original draft: J.A.M., A.O.V., H.P., P.C.C.

Writing—review & editing: J.A.M., A.O.V., H.P., S.S.-A., L.M., J.K., V.M., C.S.-G.,

J.P.-L., J.P.P., D.B., E.N., H.O., H.M., D.R.-R., P.C.-C

Project administration: A.O.V., M.L.

Funding acquisition: A.O.V., M.L., W.S., E.F.

## Data and materials availability

The authors declare that all CPAP 3054 specimens used in this study are available in the Paleontological Collection of Antarctica and Patagonia, Instituto Antártico Chileno (INACH), Punta Arenas, Chile. In addition, the authors declare that all data generated or analyzed during this study are included in this manuscript, in the Supplemental Material, and as Auxiliary files associated with this research.

## Supplementary Materials

### Supplementary Text

#### Taphonomy

Different analyses were carried out including hydraulic equivalence, bone modification and assemblage data analyses, and Voorhies groups classification. The overall study of taphonomic aspects followed traditional literature (147, 148). Only bones of duck-billed dinosaurs were recovered at the site, as is common in the fossil record of these dinosaurs, probably related to their gregarious behavior (*e.g.*, 23,27). Most of the bones were found scattered without superposition, and without any evident articulation. The size of the elements ranges between 4.8 cm and 48.1 cm, the former value corresponding to the length of the smallest caudal vertebral centrum (CPAP 5351) and the latter to the length of the largest femur (CPAP 5359). 62.5% of the elements are complete and 94% present some extent of fracture. A minimal number of three individuals (MNI) have been excavated based on the number of right femora. Relative abundance of elements from each skeletal region is shown in Table S1.

The transport of a bone element in water depends on multiple factors, such as shape, density, size and degree of articulation, in addition to the flow velocity. Disarticulated bones form distinctive classification groups (“Voorhies groups”) according to the flow speed to which they are subjected (149,150; for dinosaurs, see 151 and 152). Group 1 corresponds to bones of lower density and relatively low mass, which are affected by light currents (vertebrae and phalanges, for example); group 2 is formed by elements that are eliminated gradually, mainly due to traction; and group 3 corresponds to bones that tend to resist transport (due to their greater density and size). We found a predominance of Group 3 elements, which is consistent with the interpretation of an original deposit with the influence of low stream current transportation (see Table S2). Fluvial transport is therefore the most likely explanation for the disarticulation and scattering of the elements within the quarry.

Hydraulic equivalence allows comparing the settling velocity of fossils with that of clasts to estimate whether they were deposited at similar flow velocities, that is, by a similar process (150). Assemblages that accumulated due to fluvial processes should have a high proportion of material in hydraulic equivalence with the sediment matrix, whereas a lack of hydraulic equivalence indicates that fluvial processes were not the dominant cause of accumulation (although it does not rule out fluvial influence, 153). The average length of (complete) bones is 22 cm; large bones range between 48 cm and 32 cm and represent 35% of the complete bones. The remaining 65% range between 24.6 cm and 4.8 cm in length. When comparing these dimensions with quartz grains, using the equivalence table of (150, p. 496), most of the elements are found to be equivalent to gravel-sized grains. This means that the current velocity that transported these bones should have also been enough to transport gravel-sized grains. The matrix of the bonebed however consists predominantly of sandy mudstones with coal lenses and fine-grained sandstones. This discrepancy between the hydraulic equivalence of the bones and the matrix suggests that flow competition would have been insufficient to transport the bones a long distance, thus ruling out allochthonous assembly. This result is consistent with the interpretation based on the Voorhies group classification.

Analysis of pre-fossilization and diagenetic weathering are precluded due to current climatic conditions at the site (*i.e.*, snow cover during most of the year, temperature fluctuations and strong winds during the summer) which have an important influence on the state of preservation, accelerating weathering and erosion.

Abrasion stages can be represented by a number from 0 to 3 where 0 represents pristine fossil surfaces without signs of abrasion, and 3 corresponds to fossil bones with extremely well-rounded edges (151, 154). The elements at the bonebed show low levels of abrasion (20.6% with stage 0 and 79.4% with stage 1) that suggests minimal transport for most specimens. As already mentioned, practically all the bones are incomplete. 93.8% of the elements present fractures, most of which correspond to transverse and longitudinal fractures as caused by trampling, or contemporary exposure of the fossil material to temperature variations (climate) that produce the expansion and contraction of the material. No spiral fractures have been found in the analyzed fossils.

The different analyses above support the elements were either not transported or underwent minimal transport. This leads us to conclude that the bones are close to the original thanatocoenosis and probably to the habitat of these animals, being parauthoctonous and synchronic to some degree. The surface of many of the bones is damaged due to their exposure to current extreme weather conditions, precluding an appropriate bone modification analysis. The present data rule out death by predation and scavenging since there are no bite marks on the elements and no predator teeth have been recovered in the quarry. All lines of taphonomic evidence indicate that the reasons for the death of the individuals are biological. Considering this interpretation, the dominance of elements from Voorhies Group III is unusual. However, it may be the result of a flooding event, which would have removed elements from Voorhies Groups 1 and 2 at a time when the skeletons were exposed.

#### Reconstruction of biogeographic history

The results of analysis with BioGeoBEARS are summarized in Table S4 and Fig. S10. The best-fit model in BioGeoBEARS was the DIVALIKE + j model (AICc: 182.21), which supports a Laramidian origin for the last ancestor shared by CPAP 3054 with Hadrosauridae, and for the last ancestor shared with *Eotrachodon*, at the previous node (Fig. S10). The + j versions of all models (DEC, BAYERALIKE, and DIVALIKE) showed a better fit to the data (Table 1) and supported a Laramidian origin for the last ancestor shared by CPAP 3054 with Hadrosauridae, and for the last ancestor shared with *Eotrachodon*. The +j models support a strong role of founder effects (155), which is consistent with discrete events of dispersal across important barriers as proposed for Hadrosauroidea.

In s-DIVA, the last ancestor shared by CPAP 3054 and Hadrosauridae is recovered to have inhabited either Laramidia + Appalachia + South America, or Appalachia + South America, in equal probability. In turn, the previous last ancestor shared by CPAP 3054 with *Eotrachodon* inhabited Appalachia rather than Laramidia, as inferred by BioGeoBEARS (Fig. S11). Despite these differences, the results of s-DIVA are generally consistent with those of BioGeoBEARS in that they support the arrival of the ancestors of CPAP 3054 from North America.

Possible reasons for the differences of s-DIVA with BioGeoBEARS are that only BioGeoBEARS consider the geological age of taxa in time-calibrated trees, and that BioGeoBEARS includes a dispersal matrix that considers different dispersal probabilities between land masses.

#### Conceptual experiments with a hypothetical African taxon

When the hypothetical African taxon was placed between *Eotrachodon* and CPAP 3054, the best model (DIVALIKE + j; see Table S5) supported Laramidia + South America as the ancestral area for the last ancestor that CPAP 3054 shared with Hadrosauridae, and Laramidia as the most likely area for the last ancestor shared with *Eotrachodon* (Fig. S12).

When a hypothetical African taxon was added to our BioGeoBEARS analysis as sister taxon to CPAP 3054, the best fitting model (DIVALIKE + j; see Table S6) did not show significant changes and continued to support Laramidia as the ancestral area for both the last ancestor that CPAP 3054 shared with Hadrosauridae, and the last ancestor shared with *Eotrachodon* (Fig. S14).

When a hypothetical African taxon was added to our s-DIVA analysis in between *Eotrachodon* and CPAP 3054, this led to no changes in the areas supported for the last common ancestor shared by CPAP 3054 with Hadrosauridae, or the previous ancestor last shared by CPAP 3054 and *Eotrachodon* (Fig. S13). If the hypothetical African taxon was placed as sister of CPAP 3054, Africa is now included among poorly resolved areas for the last ancestor shared by CPAP 3054 with Hadrosauridae, but Europe and Asia remain absent, and are also absent in the immediately previous ancestor shared with *Eotrachodon*, which is still retrieved as North American (Fig. S15).

Overall, in all conceptual experiments, the results of both s-DIVA and BioGeoBEARS continued to support the arrival of CPAP 3054’s ancestors from North America.

---

**Fig. S1.**
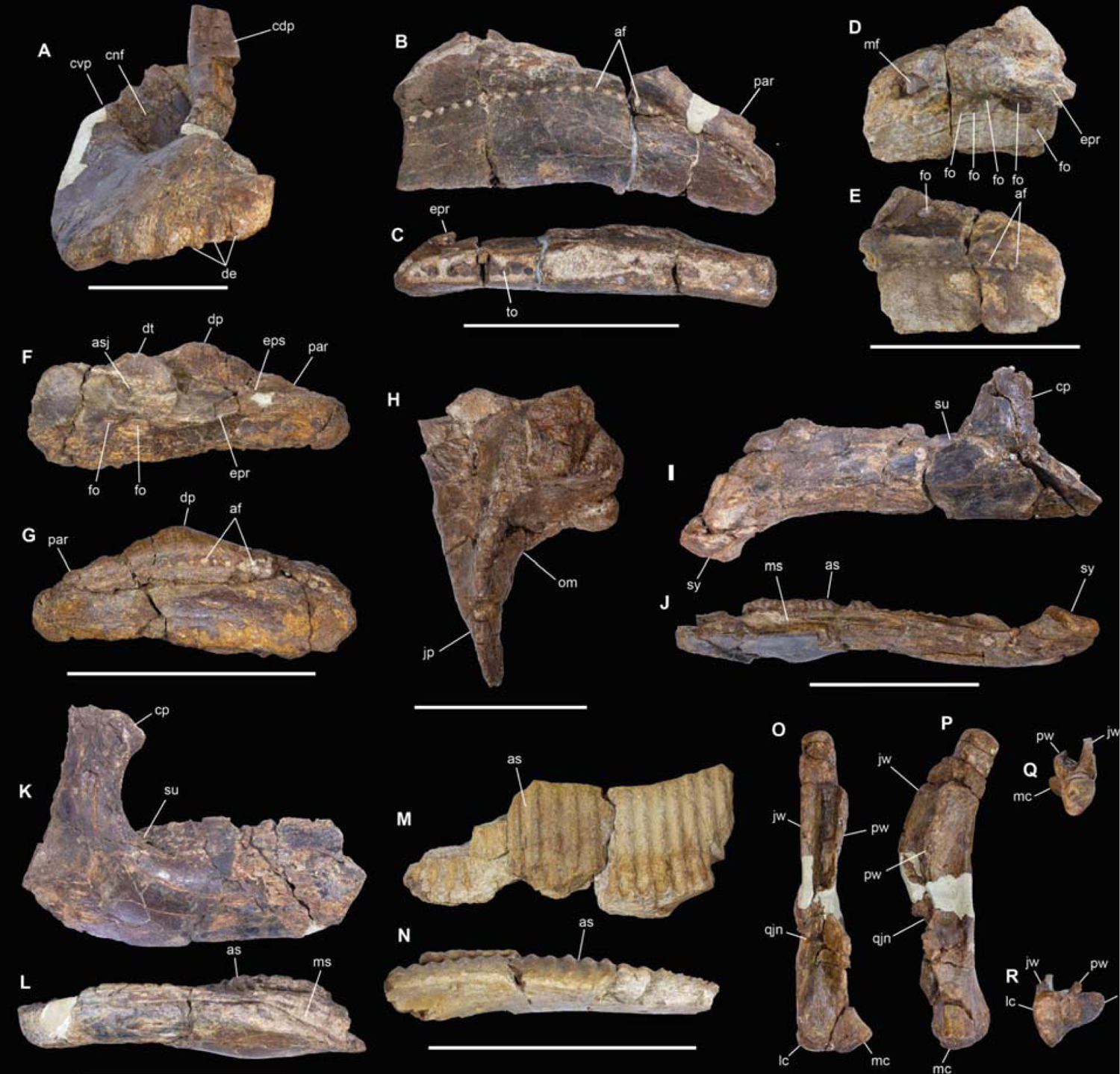
Selected cranial bones of CPAP 3054. **CPAP 5337**, right premaxilla in anterior view (**A**). **CPAP 5340**, right maxilla in lateral (**B**) and ventral (**C**) views. **CPAP 5339**, incomplete left maxilla in lateral (**D**) and medial (**E**) views. **CPAP 5338**, incomplete left maxilla of a juvenile individual in lateral (**F**) and medial (**G**) views. **CPAP 5341**, incomplete left postorbital in medial view (**H**). **CPAP 5370**, left dentary in lateral (**I**) and ventral (**J**) views. **CPAP 5342**, incomplete right dentary in lateral (**K**) and ventral (**L**) views. **CPAP 5368**, fragment of right mandibular ramus in medial (**M**) and dorsal (**N**) views. **CPAP 5343**, right quadrate in anterior (**O**), medial (**P**), proximal (**Q**) and distal (**R**) views. Abbreviations: af: alveolar foramina, af: angular facet; as: alveolar sulcus; cdp: caudodorsal process; cvp: caudoventral process; cnf: circumnarial fossa; cp: coronoid process; dp: dorsal process; dt: dorsal tubercle; d: denticles; eps: ectopterygoid shelf; epr: ectopterygoid ridge; fo: foramina; jp: jugal process; jw: jugal wing; lc: lateral condyle; ms: meckelian sulcus; mf: meckelian fossa;; mc: medial condyle; orm: orbital margin; par: palatine ridge; pw: pterygoid wing; qjn: quadratojugal notch; sy: symphysis; to: tooth. Scale bars: A, H, O-R (50 mm); B-G, I-N (100 mm).

**Fig. S2.**
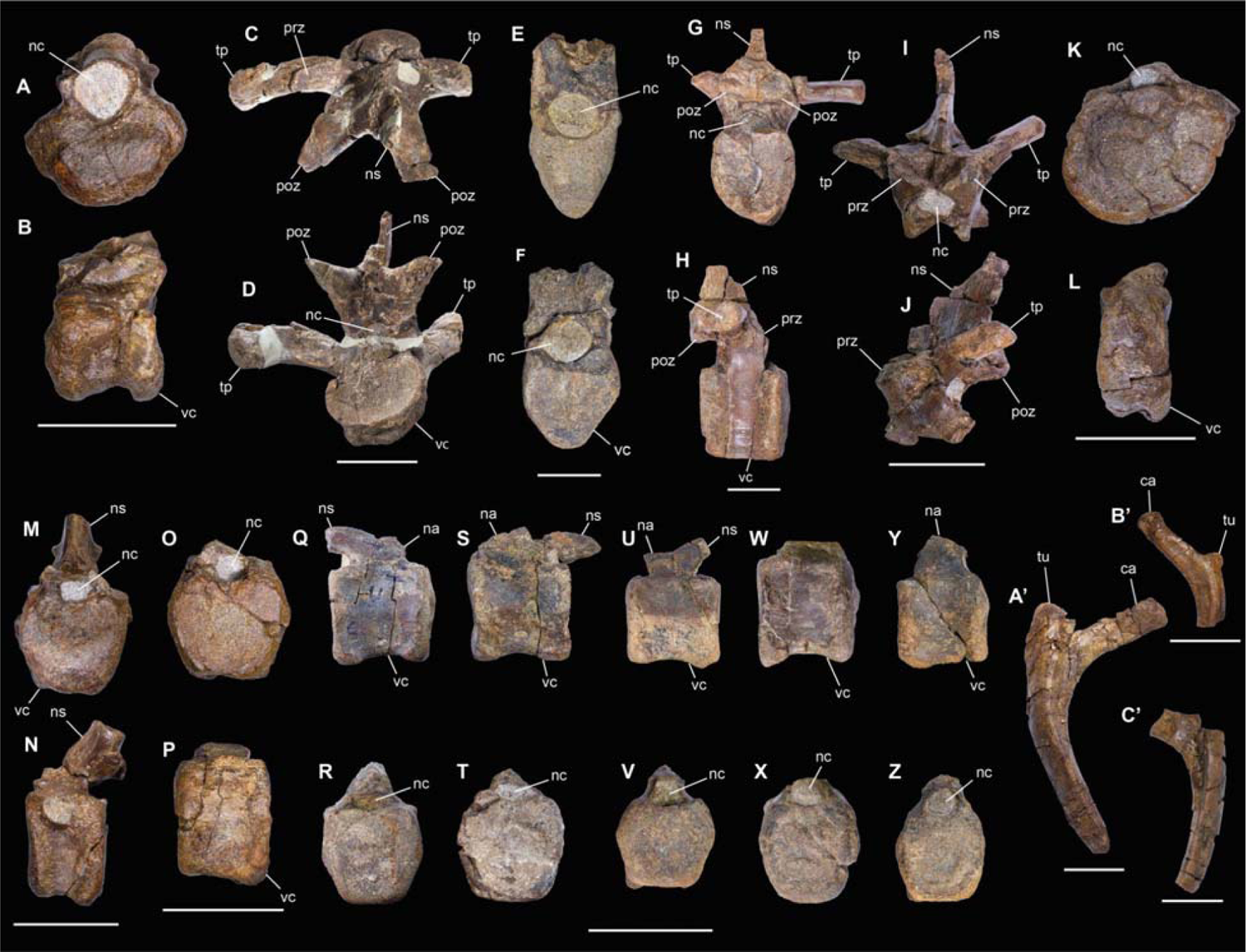
Selected axial elements of CPAP 3054. **CPAP 5380**, mid cervical centrum in anterior (**A**) and lateral (**B**) views. **CPAP 5344**, mid cervical vertebra in dorsal (**C**) and posterior (**D**) views. **CPAP 5345**, dorsal centrum in anterior (**E**) and posterior (**F**) views. **CPAP 5346**, dorsal vertebra in posterior (**G**) and lateral (**H**) views. **CPAP 5377**, dorsal neural arch in anterior (**I**) and lateral (**J**) views. **CPAP 5384**, proximal caudal centrum in anterior (**K**) and lateral (**L**) views. **CPAP 5385**, proximal caudal vertebra in anterior (**M**) and lateral (**N**) views. **CPAP 5386**, proximal caudal centrum in anterior (**O**) and lateral (**P**) views. **CPAP 5347** (**Q-R**), **CPAP 5348** (**S-T**), **CPAP 5349** (**U-V**), **CPAP 5350** (**W-X**) and **CPAP 5351** (**Y-Z**), mid**-**caudal vertebrae in lateral (**Q**, **S**, **U**, **W**, **Y**) and anterior (**P**, **S**, **U**, **W**, **Y**) views. **CPAP 5381**, right rib in anterior view (**A’**). **CPAP 5382**, proximal portion of left rib in anterior view (**B’**). **CPAP 5383**, incomplete left rib in anterior view (**C’**). Abbreviations: ca: capitulum, nc: neural canal; ns: neural spine; prz: prezygapophysis; poz: postzygapophysis; tp: transverse process; tu: tuberculum; vc: vertebral centrum. Scale bars: (50 mm).

**Fig. S3.**
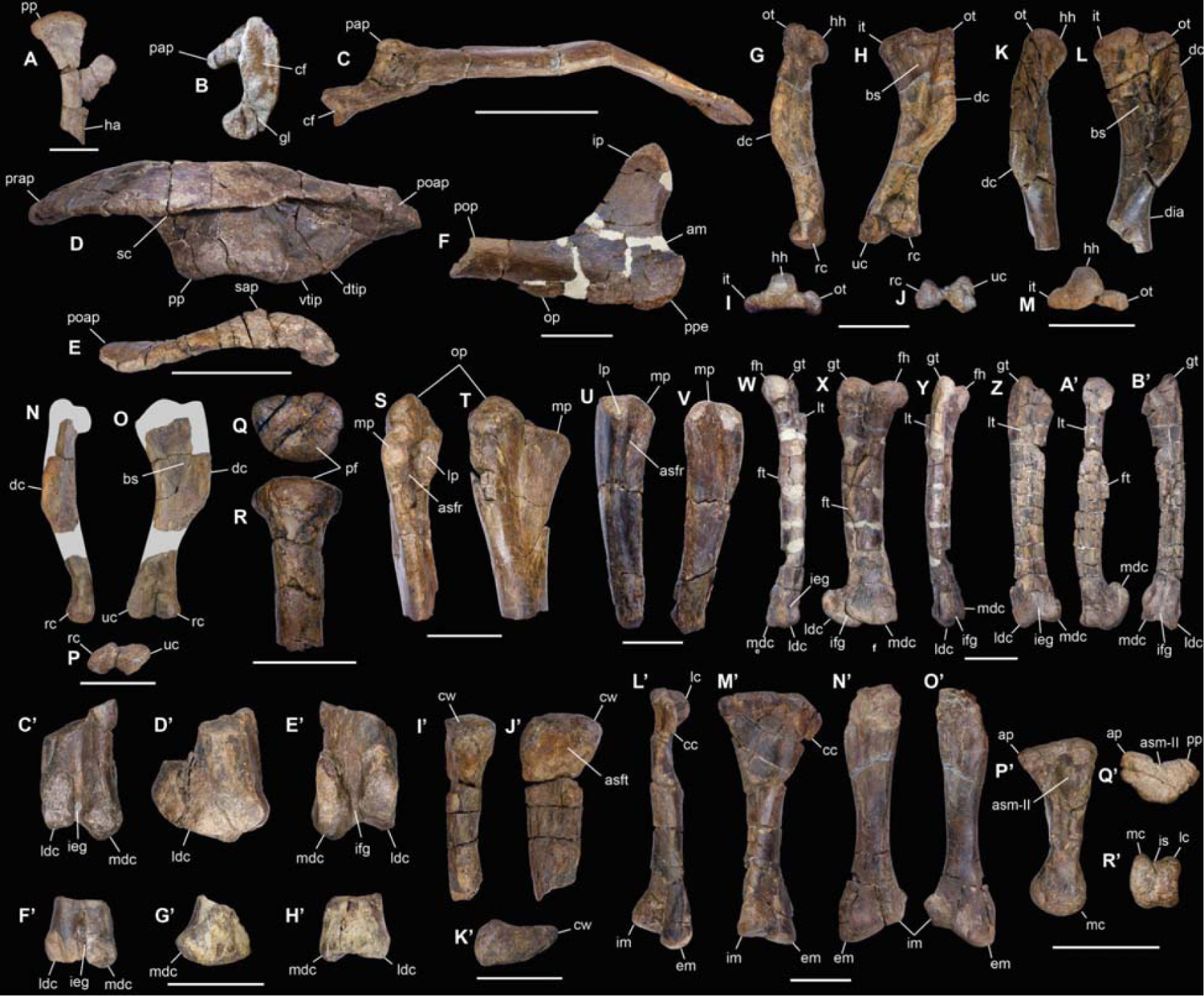
Selected appendicular elements of CPAP 3054. **CPAP 5352**, incomplete left sternum in dorsal view (**A**). **CPAP 5371**, right scapula in anterior (**B**) and dorsal (**C**) views. **CPAP 3054**, right ilium in medial view (**D**). **CPAP 5356**, postacetabular process with part of the iliac blade of a left ilium in dorsal view (**E**). **CPAP 5357**, proximal portion of right ischium in lateral (**F**) view. **CPAP 5353**, complete left humerus in ventral (**G**), anteromedial (**H**), proximal (**I**), and distal (**J**) views.**CPAP 5354**, incomplete left humerus of an immature individual in ventral (**K**), anteromedial (**L**), and proximal (**M**) views. **CPAP 5369**, incomplete left humerus in ventral (**N**), anteromedial (**O**), and distal (**P**) views. **CPAP 5379**, proximal portion of right radius in proximal (**Q**) and anterior (**R**) views. **CPAP 5355**, proximal half of a left ulna in anterior (**S**), and posteromedial (**T**) views. **CPAP 5373**, incomplete right ulna in anterior (**U**), and posteromedial (**V**) views. **CPAP 5358**, left femur in medial (**W**), posterior (**X**), and lateral (**Y**) views. **CPAP 5359**, incomplete right femur in anterior (**Z**), medial (**A’**), and posterior (**B’**) views. **CPAP 5360**, distal end of the right femur in anterior (**C’**), lateral (**D’**), and posterior (**E’**) views. **CPAP 5361**, distal portion of the right femur of a juvenile individual in anterior (**F’**), lateral (**G’**), and posterior (**H’**) views. **CPAP 5363**, proximal portion of right fibula in anterior (**I’**), medial (**J’**), and proximal (**K’**) views. **CPAP 5362**, complete left tibia in anterior (**L’**), and posterior (**M’**) views. **CPAP 5372**, incomplete left tibia in lateral (**N’**), and medial (**O’**) views. **CPAP 5364**, right metatarsal III in medial (**P’**), proximal (**Q’**), and distal (**R’**) views. Abbreviations: am: acetabular margin; asfr: articular surface for radius; asft: articular surface for the tibia; ap: anterior process; asm-III: articular surface for the metatarsal II; bs: bicipital sulcus; cf: coracoid facet; cw: cranial wing; cc: cnemial crest; dtip: dorsal tubercle of the ischial peduncle; dc: deltopectoral crest; em: external malleolus; fh: femoral head; ft: fourth trochanter; gl: glenoid; gt: great trochanter; ha: handle; hh: humeral head; ip: iliac peduncle; it: inner tubercle; ieg: intercondylar extensor groove; ifg: intercondylar flexor groove; im: internal malleolus, is: intercondylar sulcus; is: intercondylar sulcus; lp: lateral process; ldc: lateral distal condyle; lt: lesser trochanter; lc: lateral condyle; lc: lateral condyle; mp: medial process; mdc: medial distal condyle; mc: medial condyle; op: obturator process; ot: outer tubercle; op: olecranon process; pap: pseudoacromion process; pp: proximal paddle; pop: posterior process; ppe: pubic peduncle; pmc: posteromedial condyle; pp: posterior process; pp: pubic process; prap: preacetabular process; poap: postacetabular process; rc: radial condyle; sap: supraacetabular process; sc: sacral crest; su: sulcus; uc: ulnar condyle; vtip: ventral tubercle of the ischiadic peduncle. Scale bars: A, F, Q-V (50 mm); B-E, G-P, W, X, Y, Z, A’-Q’ (100 mm).

**Fig. S4.**
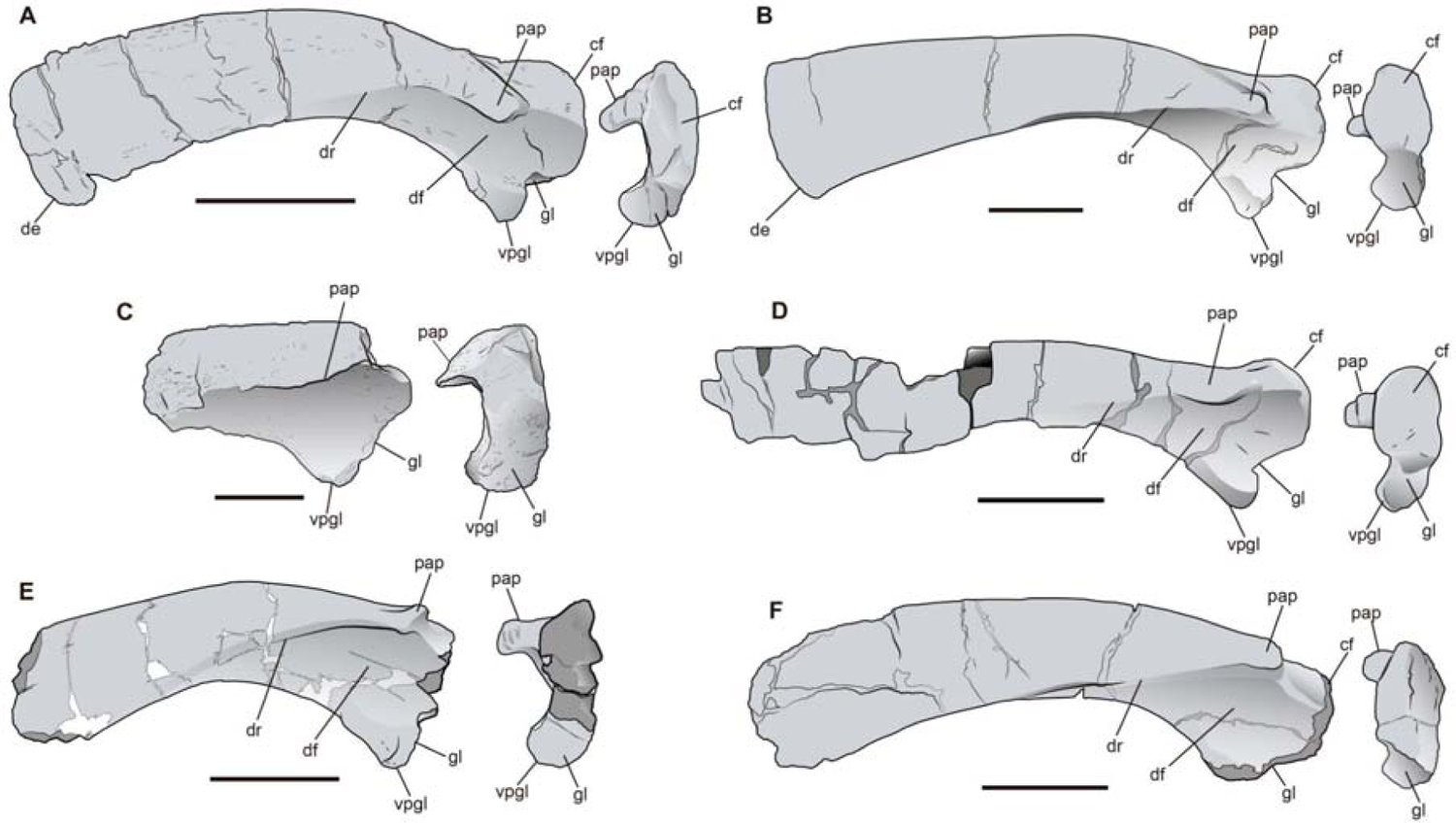
Comparison of the scapula of CPAP 3054 to that of other South American hadrosauroids. (**A**) **CPAP 5371**, right scapula in lateral and anterior views. (**B**) *Huallasaurus australis*, **MACN RN-142**, left scapula (reversed) in lateral and anterior views. (**C**) *Lapampasaurus cholinoi*, **MPHN-Pv-01**, anterior portion of left scapula (reversed) in lateral and anterior views. (**D**) *Kelumapusaura machi*, **MPCN-PV-810**, right scapula in lateral and anterior view. (**E**) *Secernosaurus koerneri*, **FMNH P13423**, right scapula in lateral and anterior views. (**F**) ‘*Willinakaqe salitrarensis*’, **MPCA-Pv SM 2**, left scapula (reversed) in lateral and anterior views. Scale bars=100 mm. Abbreviations: cf: coracoid facet; de: distal expansion; df: deltoid fossa; dr: deltoid ridge; gl: glenoid; pap: pseudoacromion process; sn: scapular neck; vpgl: ventral process of the glenoid.

**Fig. S5.**
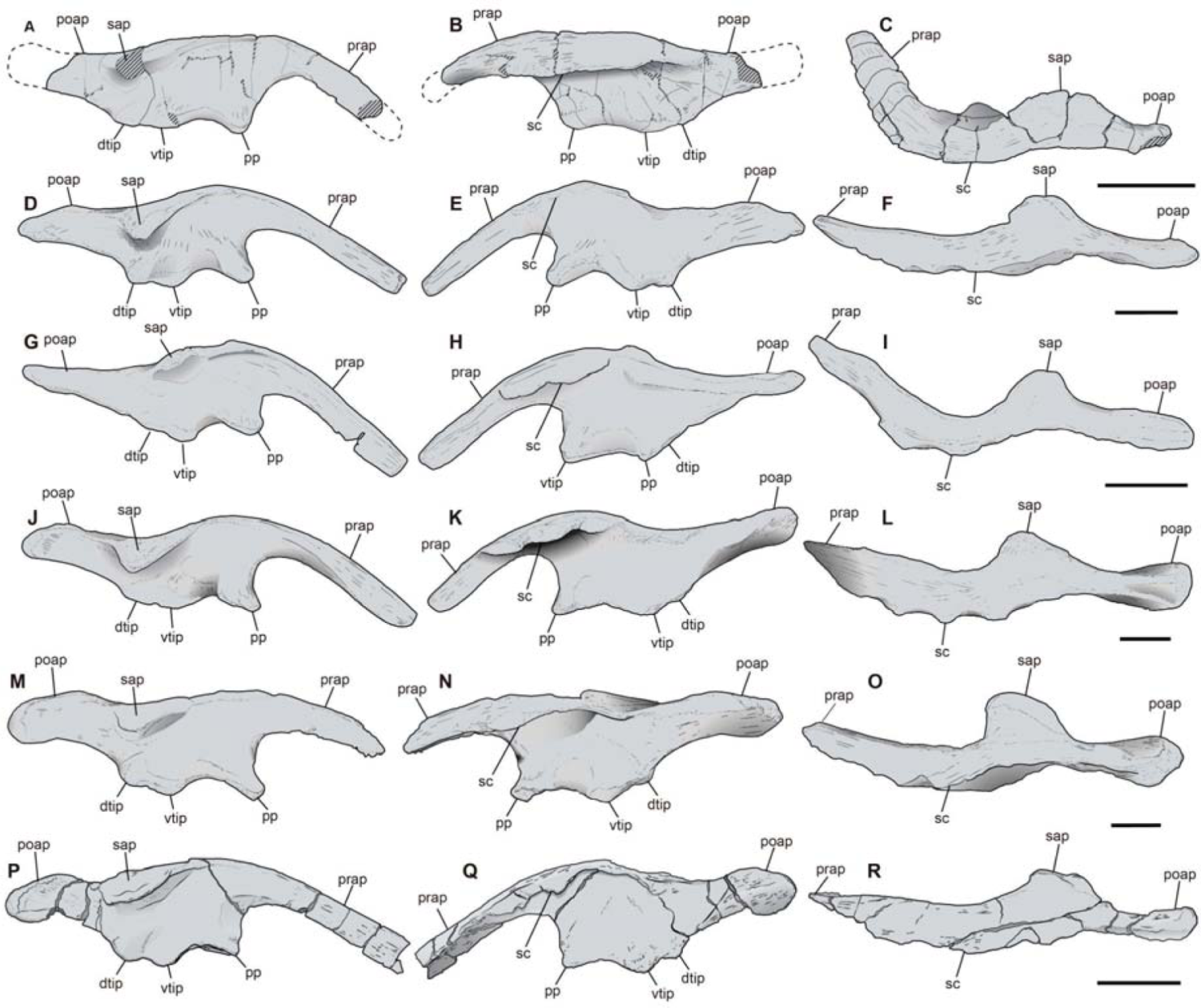
Ilia of South American duck-billed dinosaurs. **CPAP 3054** (holotype), right ilium in lateral (**A**), medial (**B**) and dorsal views (**C**). ‘*Willinikaqe salitralensis*’ **MPCA-Pv SM39**, left ilium in lateral (**D**), medial (**E**) and dorsal (**F**). *Secernosaurus koerneri*, **FMNH P13423**, right ilium in lateral (**G**), medial (**H**) and dorsal (**I**) views. *Huallasaurus australis,* **MACN-RN-02**, left ilium in lateral (**J**), medial (**K**) and dorsal (**L**) views. *Bonapartesaurus rionegrensis,* **MPCA-Pv SM2/49**, right ilium in lateral (**M**), medial (**N**) and dorsal (**O**) views. *Kelumapusaura machi*, **MPCN-PV-811**, left ilium (reversed) in lateral (**P**), medial (**Q**) and dorsal (**R**) views. Scale bars=100 mm. Abbreviations: ac: acetabulum; dtip: dorsal tubercle of the ischial peduncle; sc: sacral crest; sap: supraacetabular process; ri: ridge; sap: supraacetabular process; sc: sacral crest; pp: pubic process; prap: preacetabular process; poap: postacetabular process; vtip: ventral tubercle of the ischial peduncle.

**Fig. S6.**
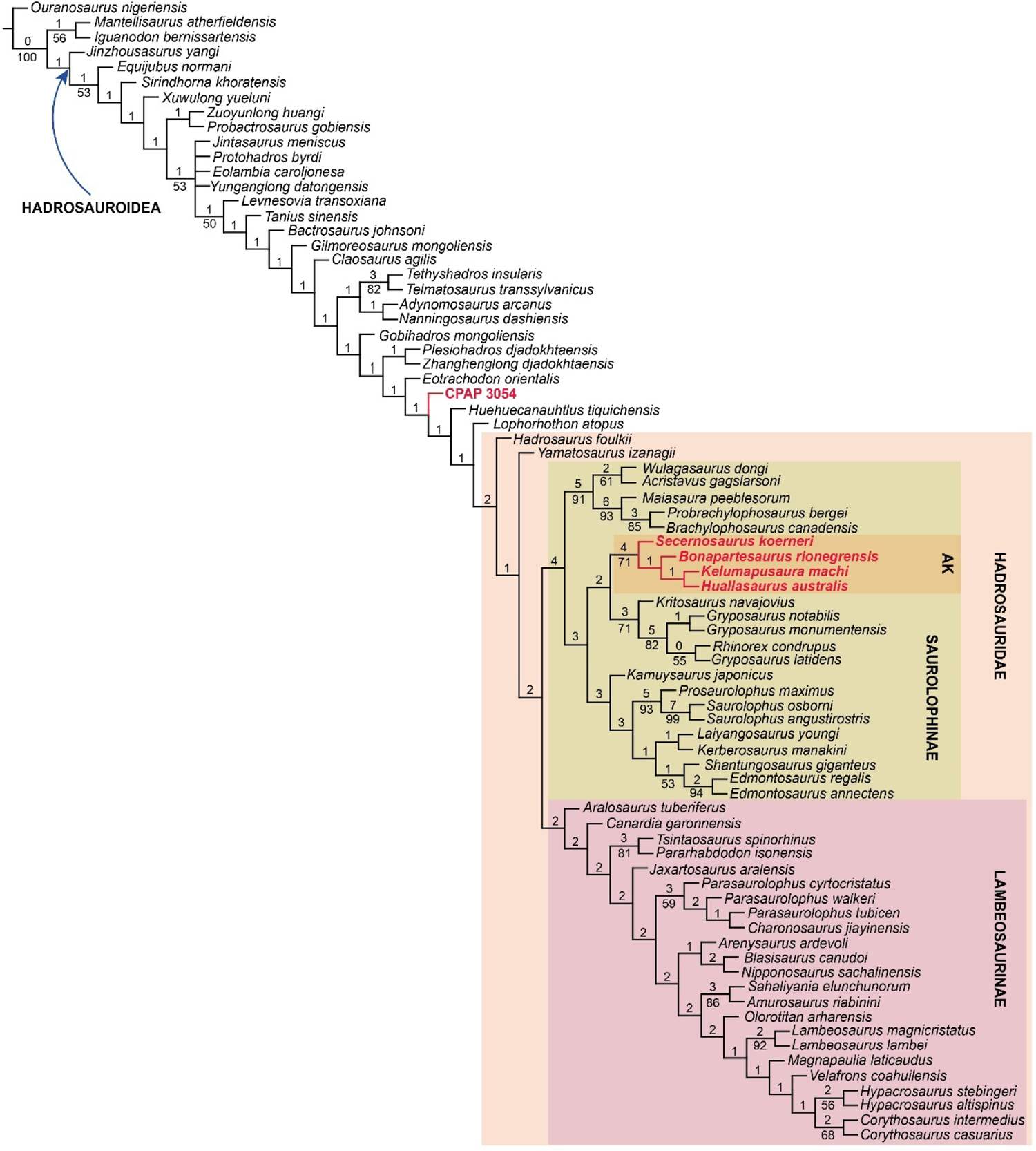
Phylogenetic analyses. Strict consensus tree (parsimony analysis) of 4 MPTs of 1312 steps (CI: 0.396; RI: 0.821). Values above nodes are Bremer support. Values beneath nodes are bootstrap proportions (5000 pseudoreplicates).

**Fig. S7.**
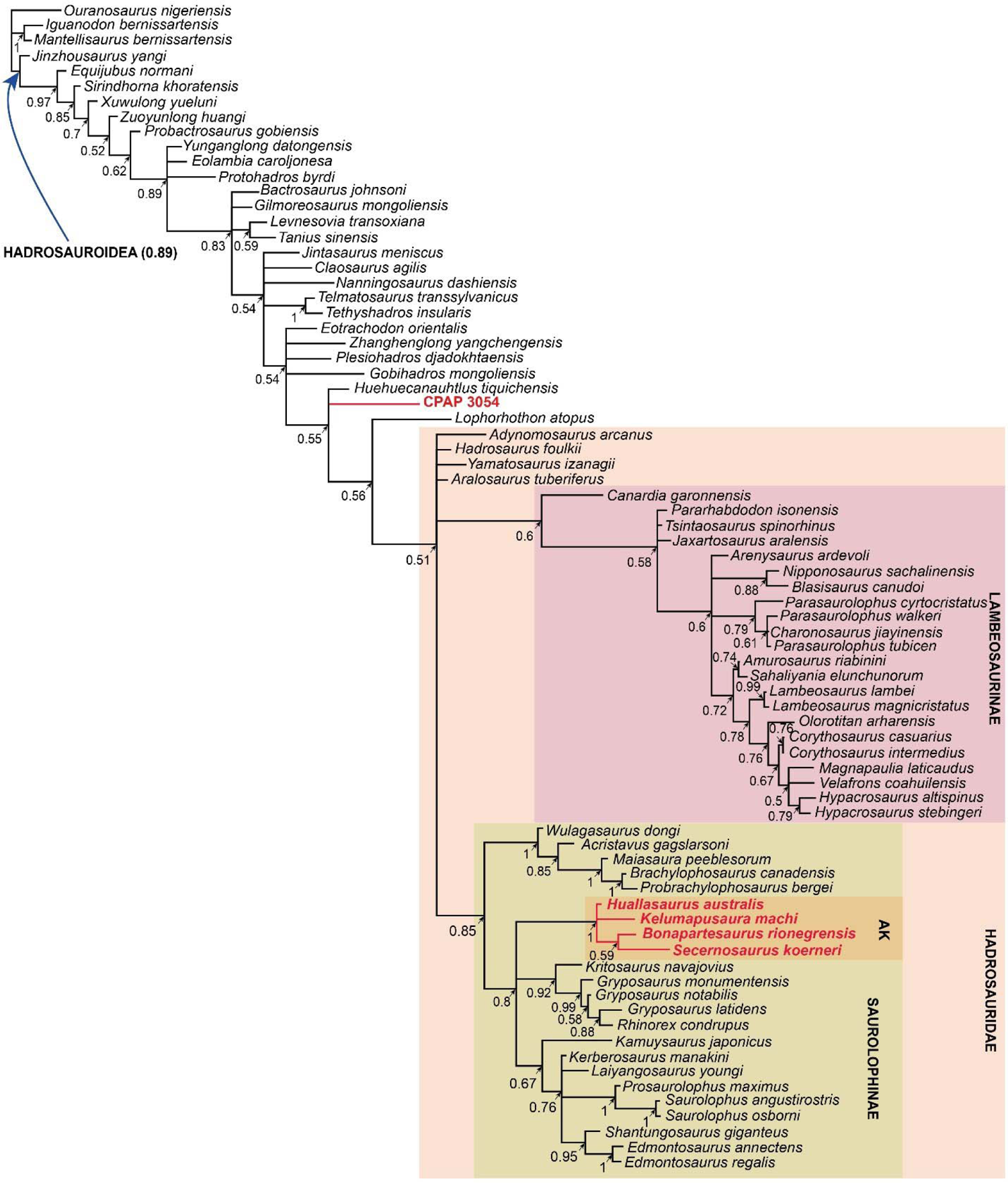
50% Majority Tree obtained by unconstrained undated Bayesian Analysis.

**Fig. S8.**
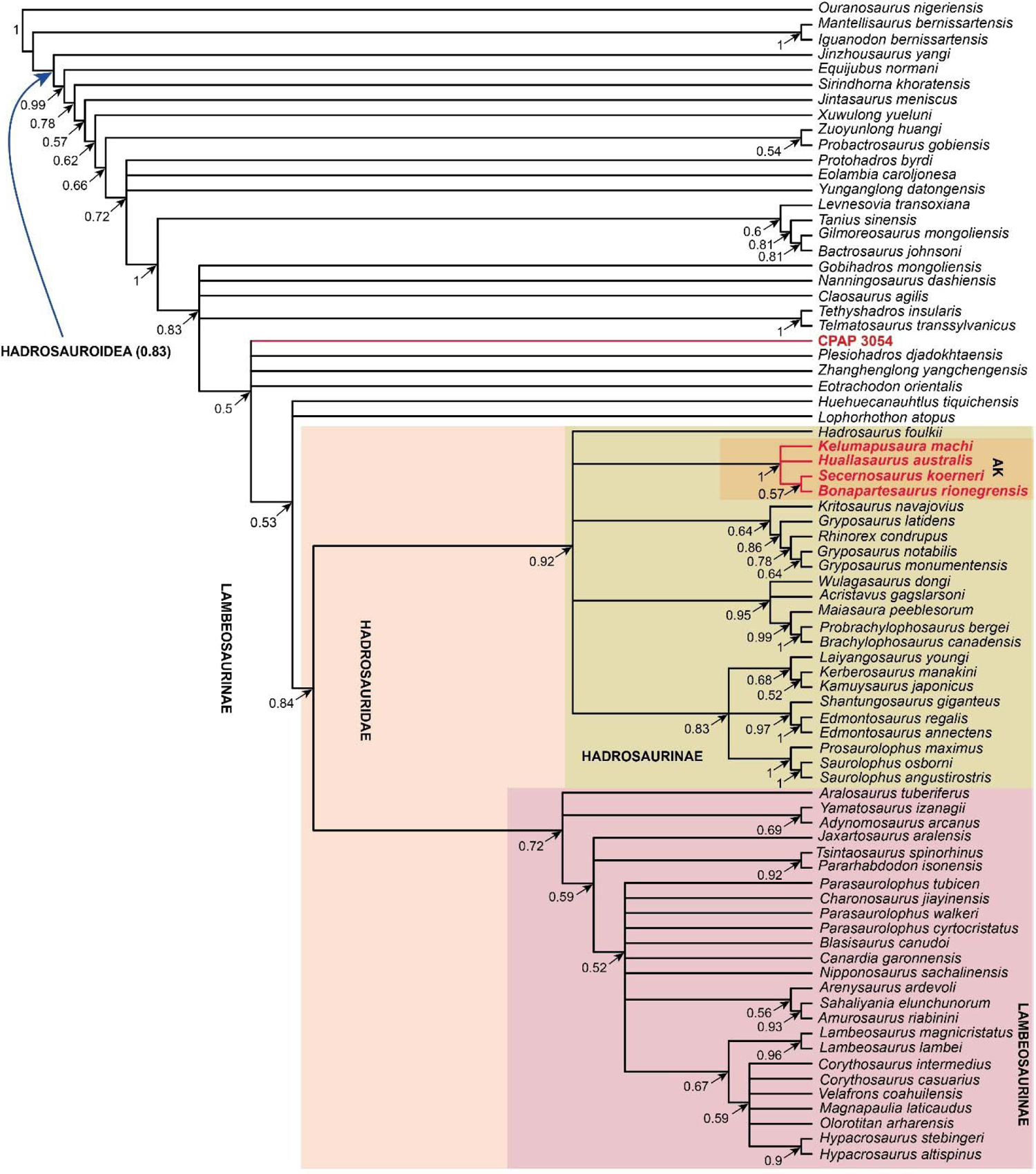
50% Majority rule tree obtained from the unconstrained tip-dated Bayesian analysis. Unlike other analyses, the clade Hadrosaurinae (Saurlophinae + *Hadrosaurus*) was recovered. Numbers above nodes correspond to the posterior probabilities. South American hadrosauroids are in blue. AK: Austrokritosauria.

**Fig. S9.**
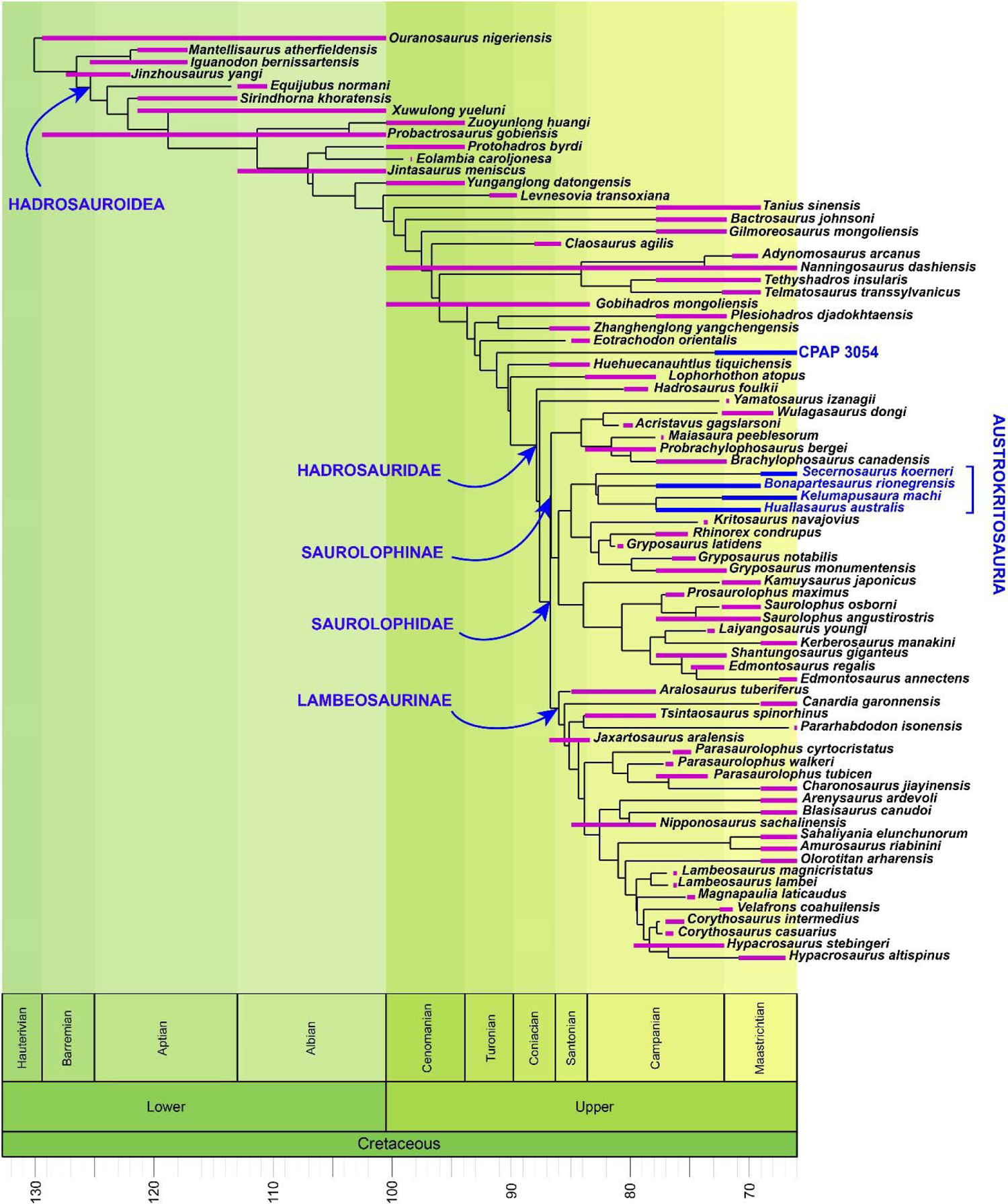
Chronostratigraphic uncertainty ranges superimposed on a time-calibrated tree obtained by parsimony analysis (see Methods for selection and time calibration of this tree). South American hadrosauroids are in blue.

**Fig. S10.**
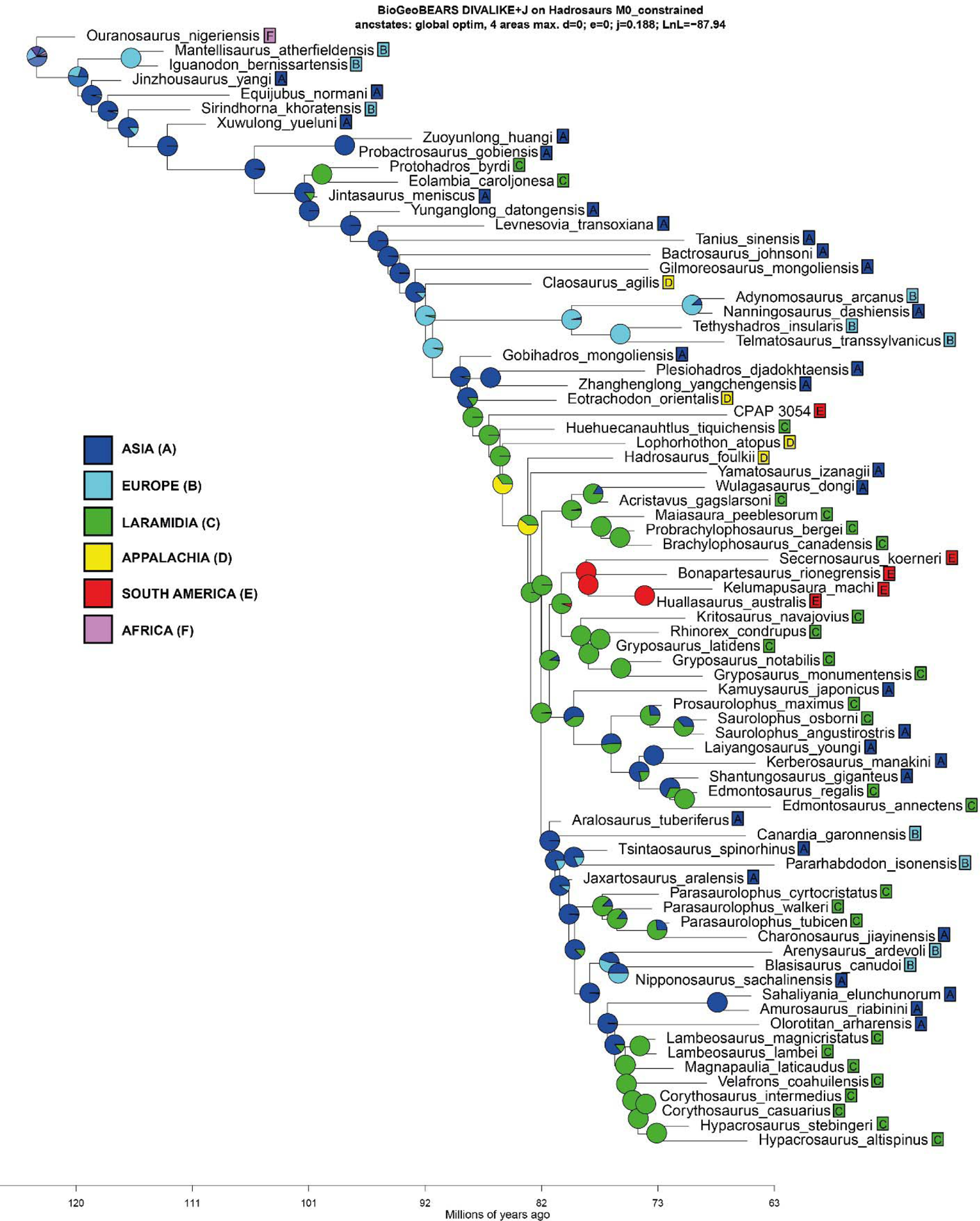
Results of Biogeographical analyses. Time-calibrated tree with BioGeoBEARS results (DIVALIKE + j model).

**Fig. S11.**
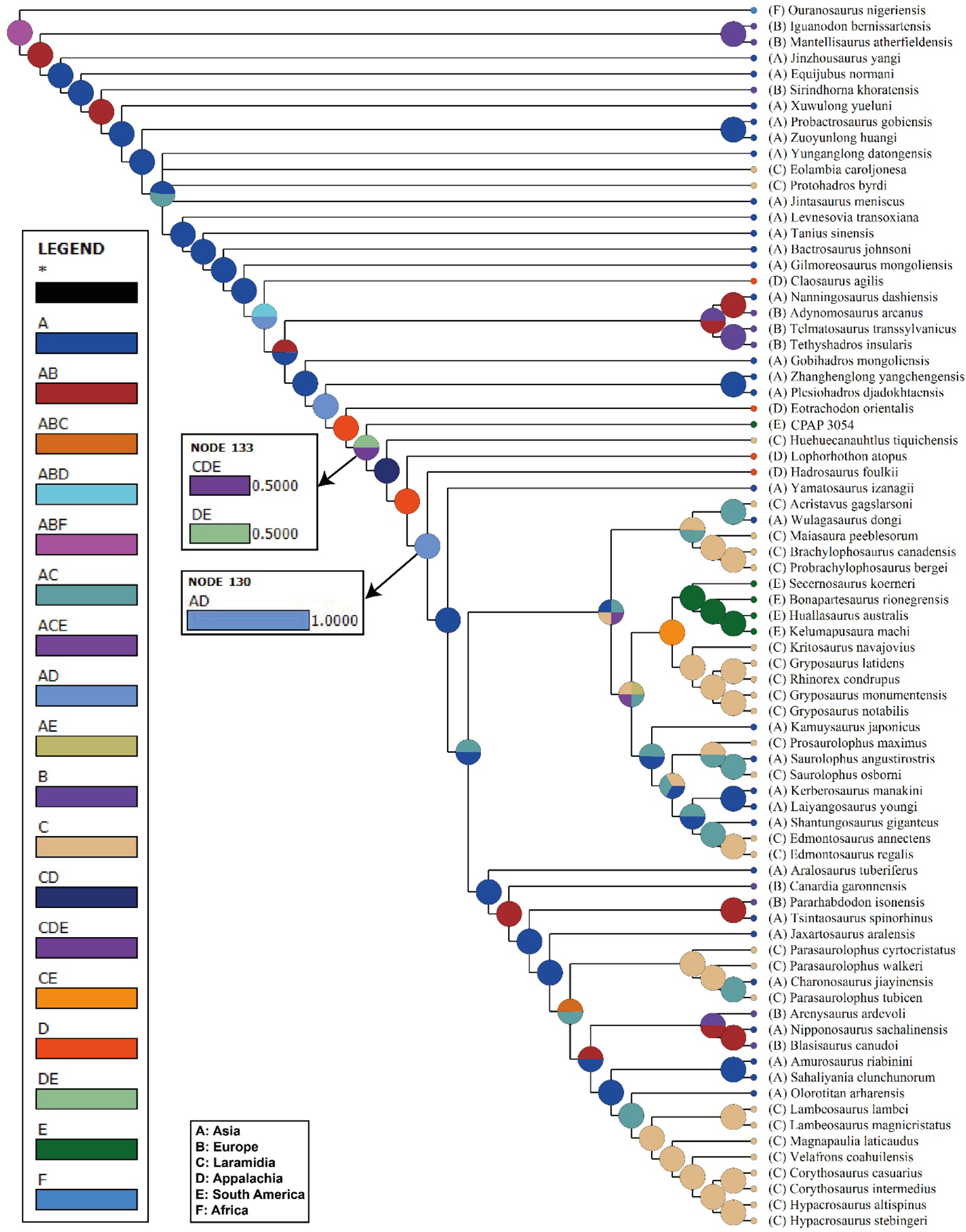
Results of Biogeographical Analyses with s-DIVA. Node 130: Hadrosauridae; Node 133: CPAP 3054 + *Huehuecanauthlus* + *Lophorhothon* + Hadrosauridae.

**Fig. S12.**
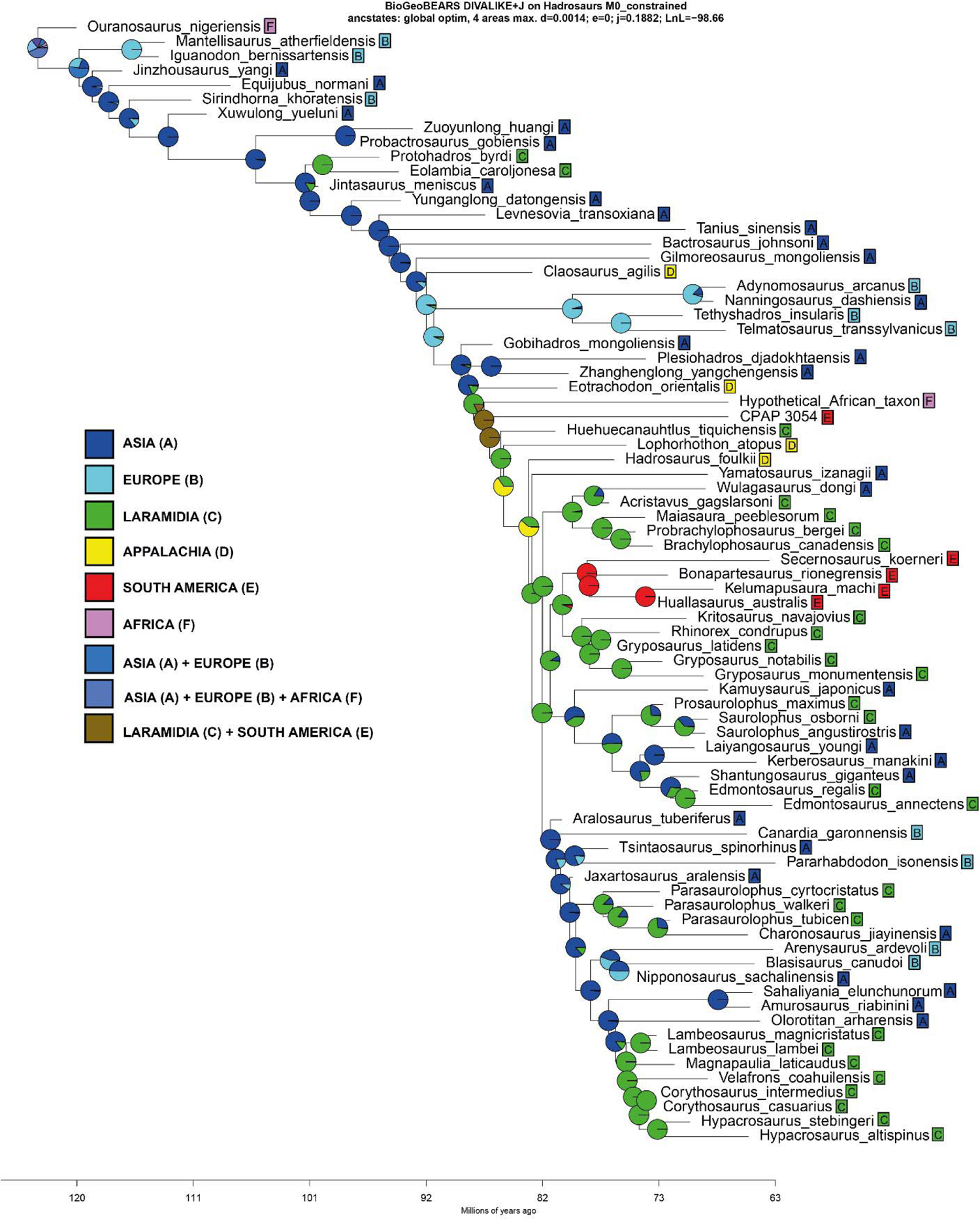
Conceptual experiments with BioGeoBEARS with a “Hypothetical African taxon” between *Eotrachodon* and CPAP 3054. Time-calibrated tree with BioGeoBEARS results (DIVALIKE + j model),

**Fig. S13.**
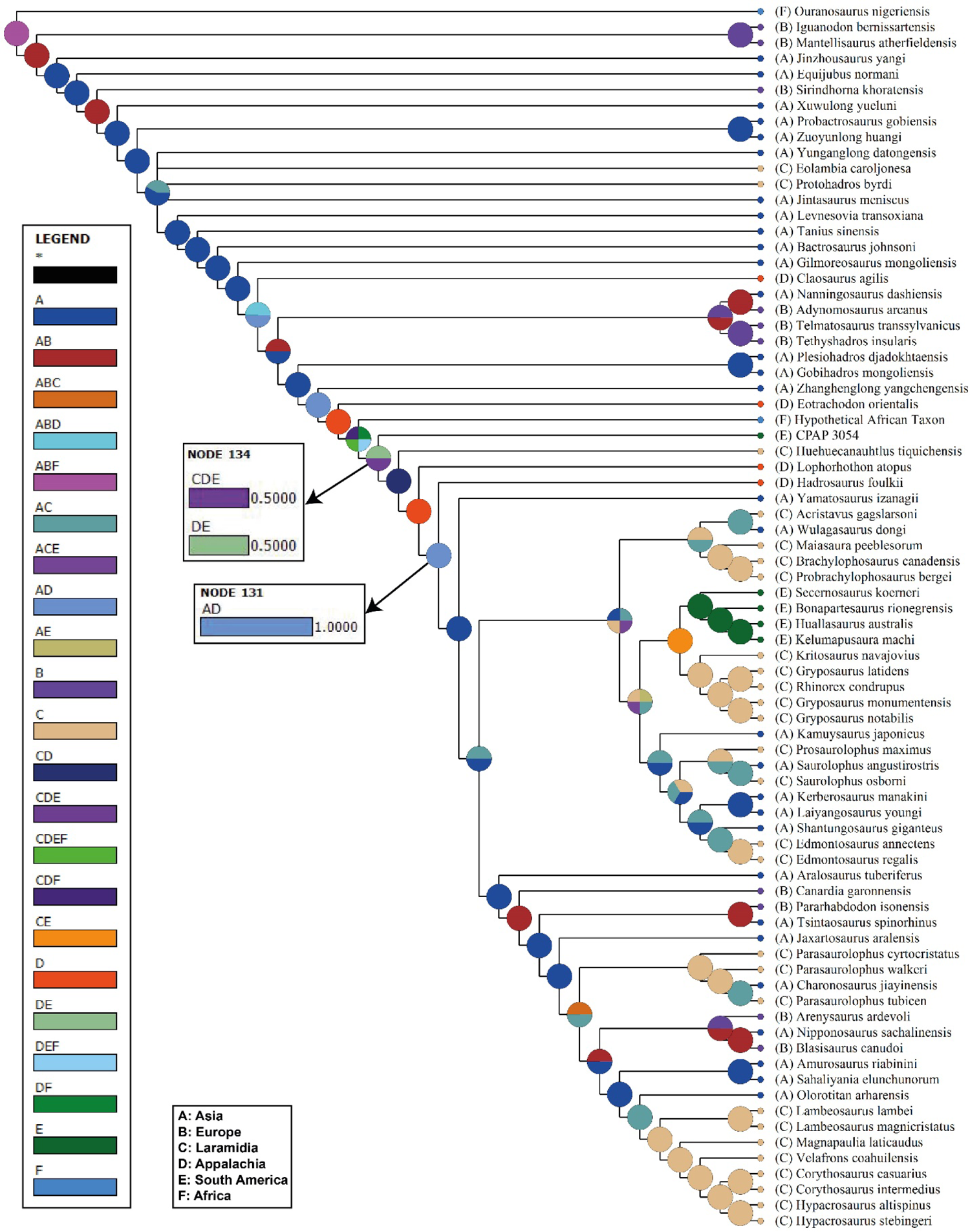
Conceptual experiment with s-DIVA with the “Hypothetical African taxon” between *Eotrachodon* and CPAP 3054. Node 131: Hadrosauridae; Node 134: CPAP 3054 + *Huehuecanauthlus* + *Lophorhothon* + Hadrosauridae.

**Fig. S14.**
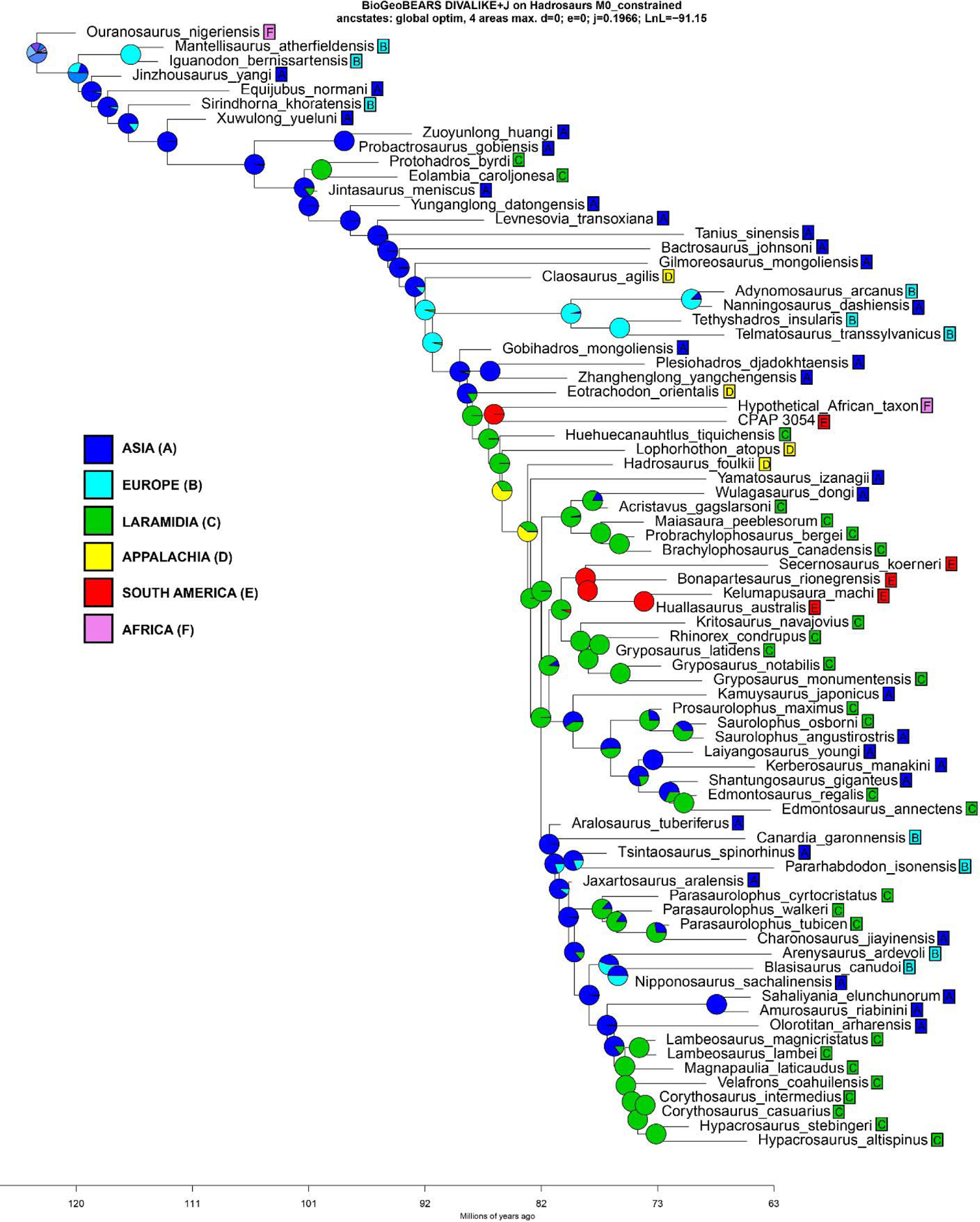
Time-calibrated tree with BioGeoBEARS results (DIVALIKE + j model), with the “Hypothetical African taxon” as sister taxon of CPAP 3054.

**Fig. S15.**
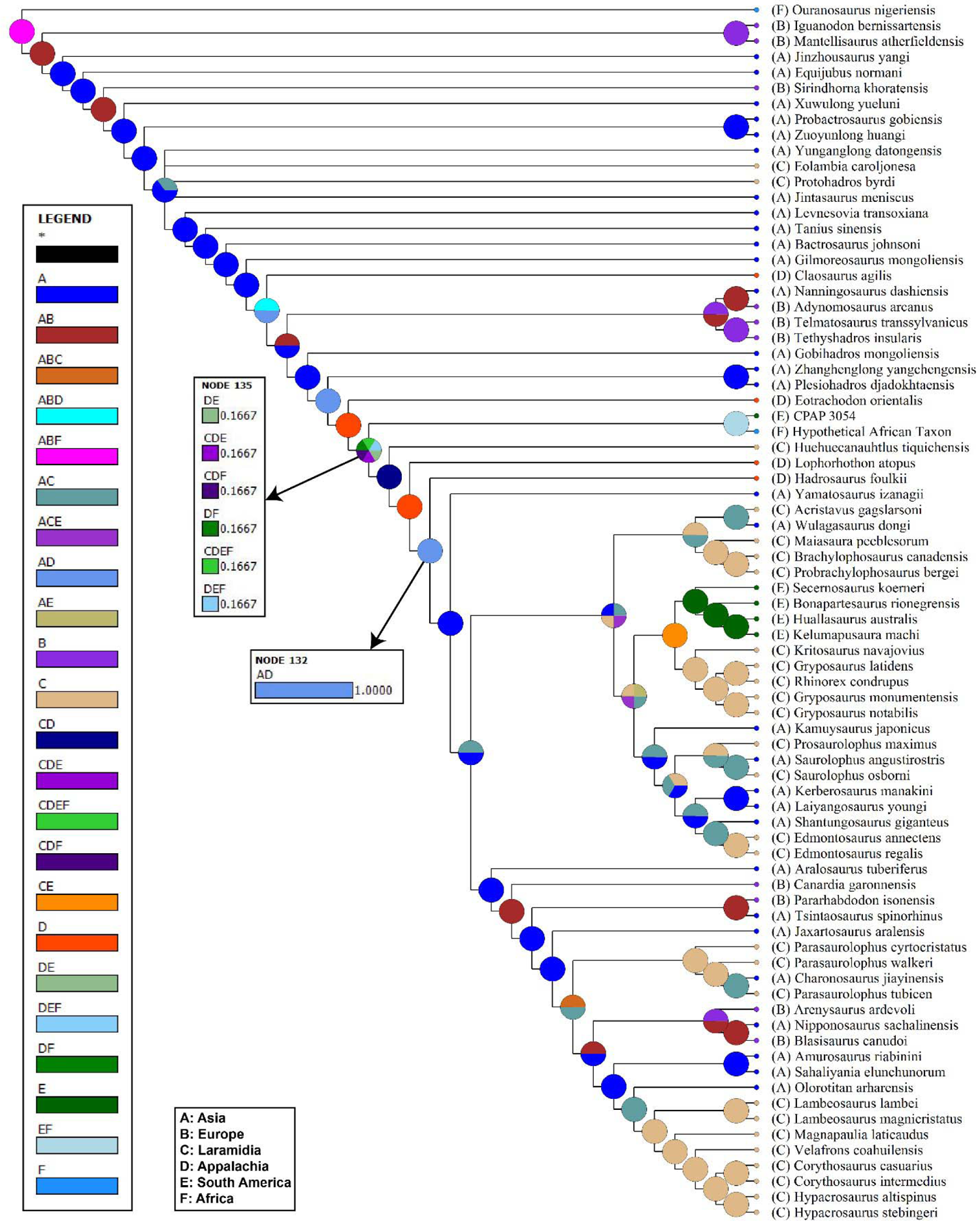
Results of s-DIVA with a hypothetical African sister taxon of CPAP 3054. Node 131: Hadrosauridae; Node 134: CPAP 3054 + *Huehuecanauthlus* + *Lophorhothon* + Hadrosauridae.

**Table S1.** Distribution of skeletal regions and series.

| Representation by skeletal region |  |  |
| --- | --- | --- |
| Region | Element representation |  |
| Skull | 9 | 26% |
| vertebrae | 8 | 24% |
| pectoral girdle | 2 | 5% |
| Forelimb | 4 | 12% |
| Pelvic girdle | 3 | 9% |
| Hindlimb | 8 | 24% |

**Table S2.** Relative frequency of elements according to Voorhies groups (based on Ryan *et al.* 2001). *Skull elements maybe over-represented because of the disarticulated skull, nevertheless it does not change the group III predominance, even if a single skull element is considered).

| Voorhies groups |  |  |  |
| --- | --- | --- | --- |
| I | Caudal vertebrae | 5 | 23.5% |
|  | Cervical vertebrae | 1 |  |
|  | Isquion | 1 |  |
|  | Metatarsal | 1 |  |
| II | Ulna | 1 | 14.7% |
|  | Fibula | 1 |  |
|  | Dorsal vertebrae | 2 |  |
|  | Sternum | 1 |  |
| III | Skull elements* | 6 | 61.8% |
|  | Dentaries | 3 |  |
|  | Humerus | 3 |  |
|  | Femur | 4 |  |
|  | Tibia | 2 |  |
|  | Scapula | 1 |  |
|  | Ilium | 2 |  |

**Table S3.**
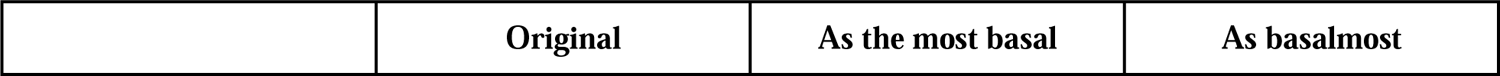
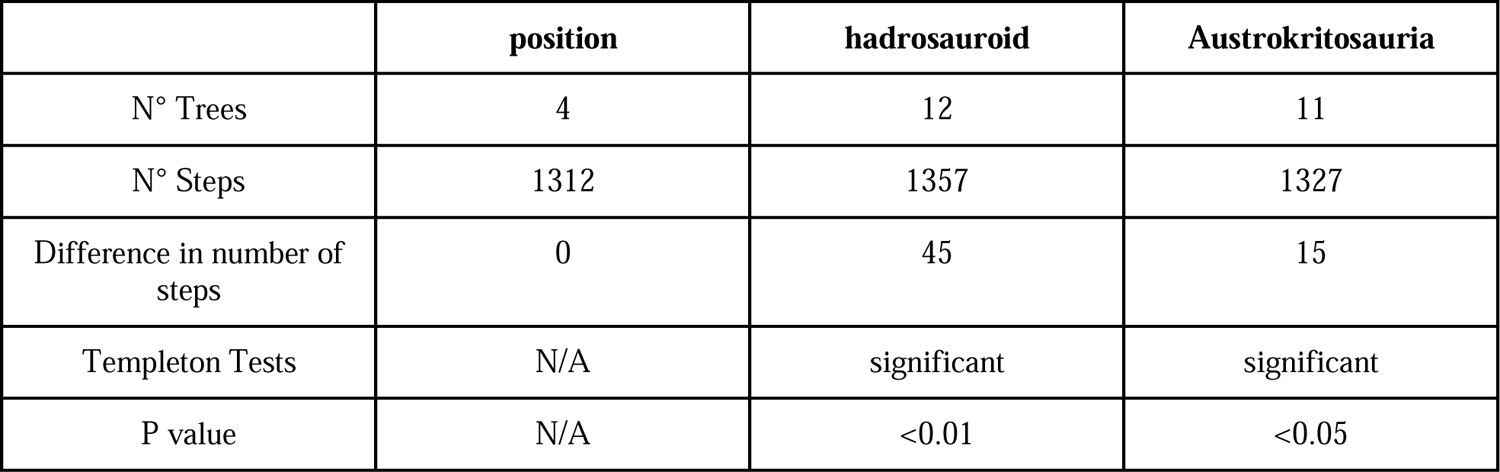
Summary of the results of phylogenetic analyses forcing CPAP 3054 into different positions. Number of trees, number of steps, and differences in steps are indicated, in addition to Templeton Tests results to assess its significance.

**Table S4.**
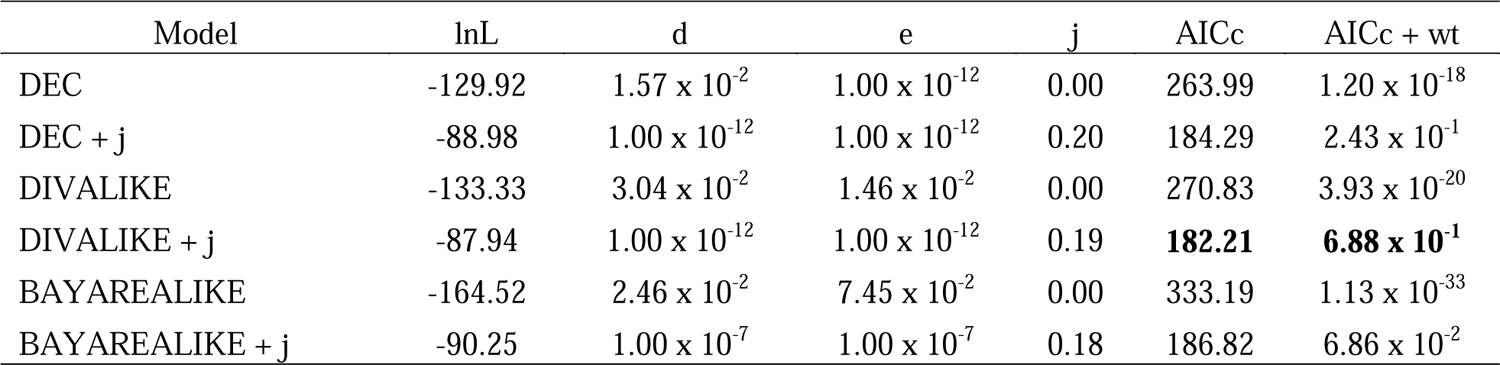
Models and parameters for each of the analyses with BioGeoBears. Values of log-Likelihood (lnL), Dispersal (d), Extinction (e), Founder effect (j), Corrected Akaike Information Criterion (AICc), and AICc Weight (AICc wt) scores from each model implemented. The best model (DIVALIKE + j) was selected based on the lowest AICc value (and higher AICc + wt).

| Model | lnL | d | e | j | AICc | AICc + wt |
| --- | --- | --- | --- | --- | --- | --- |
| DEC | -129.92 | $1.57 \times 10^{-2}$ | $1.00 \times 10^{-12}$ | 0.00 | 263.99 | $1.20 \times 10^{-18}$ |
| DEC + j | -88.98 | $1.00 \times 10^{-12}$ | $1.00 \times 10^{-12}$ | 0.20 | 184.29 | $2.43 \times 10^{-1}$ |
| DIVALIKE | -133.33 | $3.04 \times 10^{-2}$ | $1.46 \times 10^{-2}$ | 0.00 | 270.83 | $3.93 \times 10^{-20}$ |
| DIVALIKE + j | -87.94 | $1.00 \times 10^{-12}$ | $1.00 \times 10^{-12}$ | 0.19 | **182.21** | **$6.88 \times 10^{-1}$** |
| BAYAREALIKE | -164.52 | $2.46 \times 10^{-2}$ | $7.45 \times 10^{-2}$ | 0.00 | 333.19 | $1.13 \times 10^{-33}$ |
| BAYAREALIKE + j | -90.25 | $1.00 \times 10^{-7}$ | $1.00 \times 10^{-7}$ | 0.18 | 186.82 | $6.86 \times 10^{-2}$ |

**Table S5.**
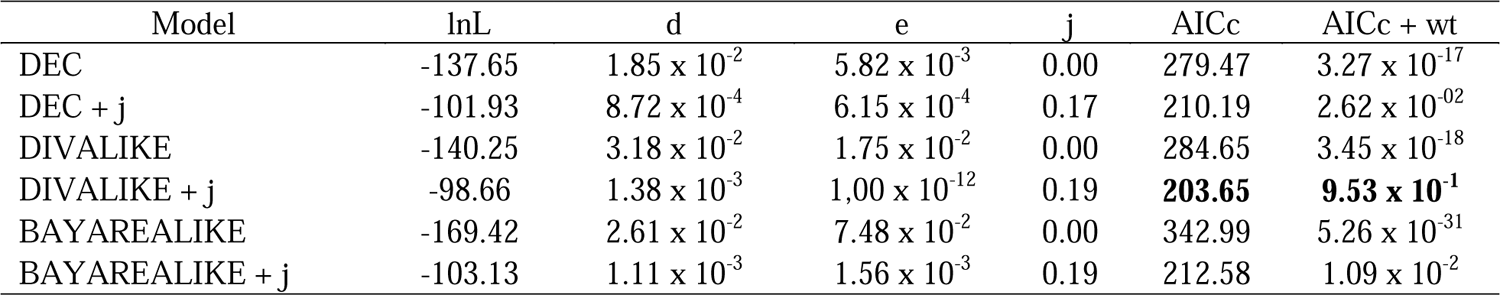
Experiment 1, “Hypothetical African Taxon between *Eotrachodon* and CPAP 3054. Models and parameters for each of the analyses with BioGeoBears. Values of log-Likelihood (lnL), Dispersal (d), Extinction (e), Founder effect (j), Corrected Akaike Information Criterion (AICc), and AICc Weight (AICc wt) scores from each model implemented. The best model (DIVALIKE + j) was selected based on the lowest AICc value (and higher AICc + wt).

| Model | lnL | d | e | j | AICc | AICc + wt |
| --- | --- | --- | --- | --- | --- | --- |
| DEC | -137.65 | $1.85 \times 10^{-2}$ | $5.82 \times 10^{-3}$ | 0.00 | 279.47 | $3.27 \times 10^{-17}$ |
| DEC + j | -101.93 | $8.72 \times 10^{-4}$ | $6.15 \times 10^{-4}$ | 0.17 | 210.19 | $2.62 \times 10^{-02}$ |
| DIVALIKE | -140.25 | $3.18 \times 10^{-2}$ | $1.75 \times 10^{-2}$ | 0.00 | 284.65 | $3.45 \times 10^{-18}$ |
| DIVALIKE + j | -98.66 | $1.38 \times 10^{-3}$ | $1.00 \times 10^{-12}$ | 0.19 | **203.65** | **$9.53 \times 10^{-1}$** |
| BAYAREALIKE | -169.42 | $2.61 \times 10^{-2}$ | $7.48 \times 10^{-2}$ | 0.00 | 342.99 | $5.26 \times 10^{-31}$ |
| BAYAREALIKE + j | -103.13 | $1.11 \times 10^{-3}$ | $1.56 \times 10^{-3}$ | 0.19 | 212.58 | $1.09 \times 10^{-2}$ |

**Table S6.** Experiment 2, “Hypothetical African taxon” as sister taxon of CPAP 3054. Models and parameters for each of the analyses with BioGeoBears. Values of log-Likelihood (lnL), Dispersal (d), Extinction (e), Founder effect (j), Corrected Akaike Information Criterion (AICc), and AICc Weight (AICc wt) scores from each model implemented. The best model (DIVALIKE + j) was selected based on the lowest AICc value (and higher AICc + wt).

| Model | lnL | d | e | j | AICc | AICc + wt |
| --- | --- | --- | --- | --- | --- | --- |
| DEC | -136.01 | $1.84 \times 10^{-2}$ | $5.61 \times 10^{-3}$ | 0.00 | 276.18 | $6.65 \times 10^{-20}$ |
| DEC + j | -92.17 | $1.00 \times 10^{-12}$ | $1.00 \times 10^{-12}$ | 0.21 | 190.67 | $2.46 \times 10^{-1}$ |
| DIVALIKE | -139.03 | $3.13 \times 10^{-2}$ | $1.47 \times 10^{-2}$ | 0.00 | 282.22 | $3.24 \times 10^{-21}$ |
| DIVALIKE + j | -91.15 | $1.00 \times 10^{-12}$ | $1.00 \times 10^{-12}$ | 0.19 | **188.63** | **$6.83 \times 10^{-1}$** |
| BAYAREALIKE | -169.47 | $2.63 \times 10^{-2}$ | $7.49 \times 10^{-2}$ | 0.00 | 343.10 | $1.95 \times 10^{-34}$ |
| BAYAREALIKE + j | -93.41 | $1.00 \times 10^{-7}$ | $1.00 \times 10^{-7}$ | 0.19 | 193.14 | $7.16 \times 10^{-2}$ |

**Table 7.** Measurements of bones of CPAP 3054. *Measurement based on preserved element.

| Specimen (CPAP) | Measured structure | Measurement (mm) |
| --- | --- | --- |
| Premaxilla (CPAP 5337) | Maximum anteroposterior length of the precircummnaral area | 98.2 |
|  | Maximum width | 86.1 |
| Maxilla (CPAP 5338) | Anteroposterior length | 136.2* |
|  | Maximum dorsoventral height | 54.8 |
| Maxilla (CPAP 5339) | Anteroposterior length | 99.1* |
| Maxilla (CPAP 5340) | Anteroposterior length | 162.1* |
| Postorbital (CPAP 5341) | Dorsoventral height | 86.1 |
|  | Anteroposterior width of the base of the jugal process | 56.3 |
| Dentary (CPAP 5342) | Anteroposterior length of the alveolar surface | 148.8* |
|  | Anteroposterior width of the alveolar sulcus | ~3.5 |
|  | Anteroposterior width of the coronoid process | 36.6 |
| Dentary (CPAP 5368) | Anteroposterior width of the alveolar sulcus | ~5.4 |
| Dentary (CPAP 5370) | Length of the proximal edentulous margin | 34.5 |
|  | Anteroposterior length of the alveolar surface | 149.3 |
|  | Length of the symphyseal process | 68.2 |
|  | Anteroposterior width of the alveolar sulcus | ~4.6 |
|  | Anteroposterior width of the coronoid process | 33.1 |
| Quadrates (CPAP 5343) | Dorsoventral height | 168.9 |
|  | Mediolateral width of the distal end | 50.5 |
| cervical centrum (CPAP 5380) | Anteroposterior length | 43.0 |
|  | Mediolateral width | 58.0 |
| cervical vertebra (CPAP 5344) | Anteroposterior length of the centrum | 53.4 |
|  | Mediolateral width of the anterior face | 53.8 |
|  | Mediolateral width of the posterior face | 57.0 |
|  | Anteroposterior length of the postzygapophyses (r/l) | 31.7*/35.5* |
| Dorsal centrum (CPAP 5345) | Anteroposterior length of the centrum | 53.9 |
|  | Dorsoventral height of the centrum | 39.5 |
| Dorsal vertebra (CPAP 5346) | Anteroposterior length of the centrum | 50.0 |
|  | Dorsoventral height of the centrum | 54.1 |
| Proximal caudal centrum (CPAP 5384) | Anteroposterior length of the centrum | 26.0 |
|  | Dorsoventral height of the centrum | 58.0 |
| Proximal caudal vertebra (CPAP 5385) | Anteroposterior length of the centrum | 37.4 |
|  | Dorsoventral height of the centrum | 56.3 |
| Mid caudal vertebra (CPAP 5347) | Anteroposterior length of the centrum | 43.3 |
|  | Dorsoventral height of the centrum | 40.5 |
| Mid caudal vertebra (CPAP 5348) | Anteroposterior length of the centrum | 41.0 |
|  | Dorsoventral height of the centrum | 43.7 |
| Mid caudal vertebra (CPAP 5349) | Anteroposterior length of the centrum | 39.3 |
|  | Dorsoventral height of the centrum | 37.6 |
| Mid caudal vertebra (CPAP 5350) | Anteroposterior length of the centrum | 42.7 |
|  | Dorsoventral height of the centrum | 44.7 |
| Sternum (CPAP 5352) | Preserved length | 137.3 |
| Scapula (CPAP 5371) | Total length | 357.1 |
|  | Maximum dorsoventral height of the proximal end | 104.3 |
|  | Dorsoventral height of the scapular blade in the midpoint | 70.7 |
|  | Dorsoventral height of the distal end | 89.7 |
| Humerus (CPAP 5353) | Total length | 306.8 |
|  | Length of the deltopectoral crest | 134.4 |
|  | Dorsoventral width of the distal end | 88.8 |
| Humerus (CPAP 5354) | Total length | 259.4* |
|  | Length of the deltopectoral crest | 129.1 |
| Humerus (CPAP 5369) | Dorsoventral width of the distal end | 78.7 |
| Ulna (CPAP 5355) | Length | 152.3 |
| Ulna (CPAP 5373) | Length | 194.2* |
| Radius (CPAP 5379) | Mediolateral width of the proximal end | 47.0 |
| Ilium (CPAP 3054) | Length of the preacetabular process | 134.2* |
|  | Height of the base of the preacetabular process | 5.7 |
|  | Anteroposterior length of the iliac blade | 150.3 |
|  | Height of the iliac blade between the dorsal surface and the pubic process | 96.4 |
|  | Anteroposterior length of the supraacetabular process | 124.1 |
| Incomplete ilium (CPAP 5356) | Anteroposterior length of the supraacetabular process | 96.3 |
|  | anteroposterior length of the postacetabular process | 97.8 |
| Ischium (CPAP 5357) | Height of the iliac peduncle | 67.1 |
|  | Height of the pubic peduncle | 45.1 |
| Femur (CPAP 5358) | Total length | 485.1 |
|  | Mediolateral width of the distal end | 70.5 |
| Femur (CPAP 5359) | Total length | 475.2 |
|  | Mediolateral width of the distal end | 70.6 |
| Femur (CPAP 5360) | Mediolateral width of the distal end | 84.8 |
| Femur (CPAP 5361) | Mediolateral width of the distal end | 74.3 |
| Tibia (CPAP 5362) | Total length | 437 |
|  | Length of the cnemial crest | 164.9 |
| Tibia (CPAP 5372) | Total length | 447.1* |
| Fibula (CPAP 5363) | Anteroposterior length of the proximal end | 53.4* |
| Metatarsal III (CPAP 5364) | Length | 169.2 |
|  | Mediolateral width of the shaft | 41.4 |

